# An RNA ligase shapes transcriptional profiles, neural function, and behaviour in the developing larval zebrafish

**DOI:** 10.64898/2025.12.01.691575

**Authors:** Fiona S. Klusmann, Anna C. Kögler, Katja Slangewal, Onur Önder, Heike Naumann, Andreas Marx, Armin Bahl, Patrick Müller

## Abstract

RNA ligases are essential for the repair, splicing, and editing of RNA across various biological systems. Recently, a new enzyme that catalyses 5’-3’ RNA ligation – RNA ligase 1 (Rlig1) – was identified *in vitro*. However, the *in vivo* biological functions of Rlig1 have remained elusive. Here, we reveal the role of Rlig1 during vertebrate development using embryonic and larval zebrafish as a model system. We found that *rlig1* mRNA is maternally deposited and present ubiquitously during early embryogenesis, whereas at larval stages it localises to the brain and eyes. Interestingly, CRISPR/Cas9-generated *rlig1* knockout zebrafish exhibited no overt morphological abnormalities, but showed reduced behavioural responsiveness to visual stimuli along with massively perturbed transcriptomes and widespread dysregulation of core metabolic and translational pathways. Brain-wide calcium imaging in *rlig1* knockout larvae revealed decreased neuronal activity in key regions for visual processing, consistent with the observed behavioural defects. Together, our findings identify a role for Rlig1 in maintaining the integrity and function of the nervous system and uncover a new link between neuronal RNA processing, development, and sensory-motor computation.

## Introduction

RNA ligases are ubiquitous across all forms of life^1–4^. These enzymes catalyse the formation of phosphodiester bonds between RNA molecules. In doing so, they play essential roles in repairing, splicing, and editing RNAs^2^. RNA ligases are best known for their roles in the repair and splicing of tRNAs. For example, during *T4* bacteriophage infection in *E. coli*, host tRNAs are damaged, which can be counteracted by the activity of an *E. coli*-encoded RNA ligase^5^. Moreover, RNA ligases contribute to the final step of the splicing reaction of intron-containing tRNAs across all phylogenetic domains^3^. In addition, members of the Rnl2 family are involved in the RNA-guided editing of mitochondrial pre-mRNAs in trypanosomes^3,6^. These examples highlight the broad functional scope of RNA ligases and underscore their biological significance across diverse organisms.

To date, only two RNA ligases have been identified in vertebrates: the RNA-splicing ligase RtcB homolog (RtcB) and RNA ligase 1 (Rlig1). RtcB exhibits 3’-5’ ligase activity and is involved in the splicing of specific intron-containing tRNAs^7,8^. During this process, tRNA splicing endonucleases generate one exon with a 2′,3′-cyclic phosphate and one with a 5′-hydroxyl group^3^. RtcB catalyses the ligation of such RNA fragments by joining the two ends^3,7^. In addition to its role in tRNA processing, RtcB is also required for the unconventional splicing of *xbp1* mRNA, which encodes a key transcription factor in the unfolded protein response^7,9^. In contrast, the recently identified RNA ligase 1 (Rlig1) catalyses 5′-3′ ligation by joining RNA fragments with a 5′-phosphate and a 3′-hydroxyl group^4^.

Structural analysis of Rlig1 revealed the presence of a nucleotidyltransferase (NT) domain that harbours several conserved ligase-specific sequence motifs, including motifs I, Ia, III, IV, and V (Figure 1a)^4^. Notably, Rlig1 contains a lysine at position 57 within the conserved KX(D/H/N)G motif (Figure 1b), a signature of the nucleotidyltransferase superfamily, which is essential for adenylating the RNA 5′-phosphate^10^. Substrate studies *in vitro* revealed a preference of human Rlig1 to catalyse the ligation of RNA-dumbbell structures that possess a nicked loop region^4^. Further investigation implicated Rlig1 in maintaining ribosomal RNA integrity, since a cultured human cell line defective for Rlig1 showed increased rRNA degradation upon oxidative stress^4^. In contrast, lower tRNA levels in the brains of *rlig1* knockout mice suggested tRNAs as potential targets of Rlig1^11^. Further studies on Rlig1 have identified inhibitors of its enzymatic activity^12^ as well as small molecules that exhibit synthetic lethality upon *rlig1* knockout in human cells^13^. More recently, Rlig1 has also been implicated in noncanonical circRNA biogenesis during herpesvirus infection, highlighting its emerging versatility in RNA metabolism^14^. However, the *in vivo* function of Rlig1 in multicellular organisms remains elusive.

**Figure 1.**
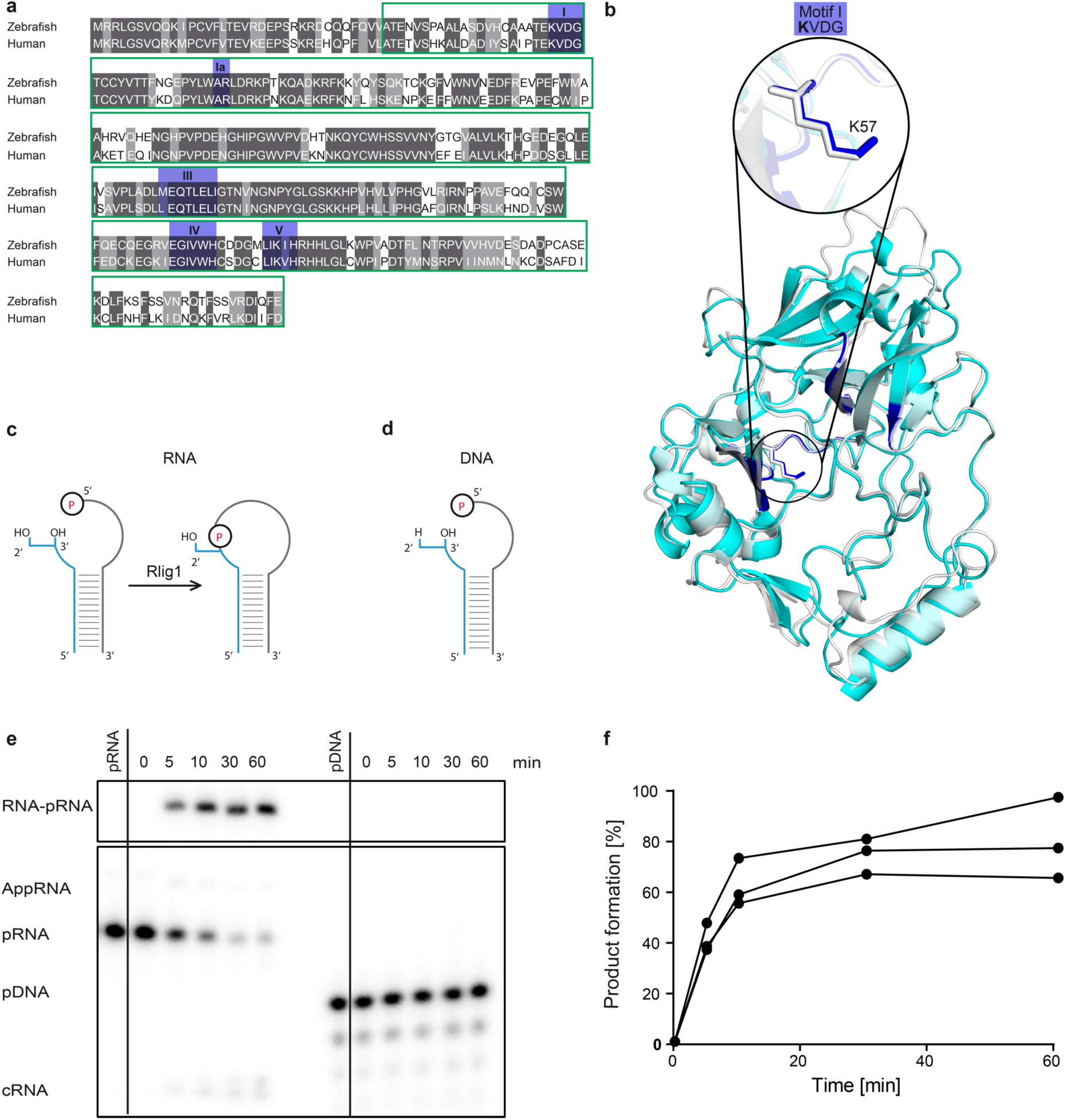
Zebrafish Rlig1 exhibits 5’-3’ single-stranded RNA ligase activity. **(a)** Protein sequence alignment of zebrafish Rlig1 and human Rlig1. The nucleotidyltransferase domain is boxed in green, and motifs I, Ia, III, IV, and V are shaded blue. **(b)** Structural alignment of AlphaFold predictions of human Rlig1 (grey) and zebrafish Rlig1 (cyan)^77–79^. The motifs I, Ia, III, IV, and V of the nucleotidyltransferase domain are depicted in blue, as in (a). The catalytic lysine residue (K57) within motif I is highlighted. **(c,d)** Schematic depiction of an RNA (c) and DNA (d) hairpin substrate consisting of two oligonucleotide strands. The unlabelled strand is depicted in blue and bears the free 3’-hydroxyl group. The ³²P-labeled 5’-end of the grey strand is depicted in red. Rlig1 catalyses the ligation of the open loop in the RNA hairpin substrate but is unable to ligate the DNA hairpin substrate, as shown in panel (e). **(e)** Time-dependent analysis of Rlig1 ligation activity. Reactions were performed for 0, 5, 10, 30, and 60 min using the RNA hairpin (depicted in c) and the DNA hairpin (depicted in d) as substrates. Phosphorimaging of the resolved radioactively labelled oligonucleotides shows that Rlig1 ligates the RNA substrate, whereas the DNA substrate remains unligated. Representative image from n = 3 independent experiments is shown. cRNA refers to circularised RNA; pDNA and pRNA denote phosphorylated (labelled) DNA or RNA, respectively; AppRNA represents the RNA-adenylate intermediate, and RNA-pRNA indicates the ligated RNA product. **(f)** Quantification of RNA ligation over time. The formation of the ligated RNA hairpin product is shown as a function of time. Data represent n = 3 independent experiments.

Here, we provide evidence of the cellular role and functional relevance of Rlig1 in zebrafish (*Danio rerio*), which possesses a single Rlig1 homolog. Using biochemical analyses, we found that zebrafish Rlig1 has similar ligation activity and substrate specificity as the previously characterised human enzyme. We then generated CRISPR/Cas9-mediated *rlig1* knockout mutants and applied a multi-level experimental approach to characterise the developmental and physiological consequences of Rlig1 loss *in vivo*, including spatiotemporal expression analysis of *rlig1* mRNA during embryogenesis and neurodevelopment, behavioural assessment of visual motion responsiveness, calcium imaging of neuronal activity, and transcriptome profiling. We found that *rlig1* transcripts are maternally deposited and later enriched in the brain and eyes. Loss of Rlig1 led to reduced responsiveness to visual motion cues, accompanied by diminished neuronal activity in visual processing centres. Transcriptomic analyses revealed widespread, stage-specific gene expression changes and consistent enrichment of pathways related to oxidative phosphorylation, ribosome function, and neurotransmitter signalling. Through this integrative analysis, we uncovered the physiological relevance of Rlig1 and identified affected cellular pathways. Our results provide novel insights into RNA ligase function during early vertebrate development and suggest that Rlig1 plays an important role in the building and maintenance of functional neural networks for signal processing and behavioural regulation.

## Results

### RNA substrate selectivity and activity of recombinant zebrafish Rlig1

To determine whether the zebrafish Rlig1 homolog has 5’-3’ RNA ligation activity similar to human Rlig1^4^, we expressed recombinant full-length zebrafish Rlig1 in *E. coli* and purified the protein by affinity and anion exchange chromatography. The ligation activity and selectivity of zebrafish Rlig1 for potential RNA and DNA substrates was assessed using hairpin constructs, as broken RNA hairpins have been found to be preferential substrates of human Rlig1 due to their loop structure^4^. These constructs consist of a 5’ strand of 10 nucleotides (nt) bearing a free 3’ hydroxyl group and a 17 nt 3’ strand labelled with ^32^P-phosphate at its 5’ terminus (blue and grey, respectively, in Figure 1c–d). The two oligonucleotide strands anneal, forming a hairpin with a nick in the loop region. Successful ligation yields a 27 nt hairpin, which migrates distinctly from the unligated educt in a denaturing polyacrylamide gel^4^. To initiate the ligation reaction, we incubated both oligonucleotide strands with zebrafish Rlig1 in the presence of ATP. We monitored product formation of the RNA and DNA constructs over a period of 1 h (Figure 1e). The formation of the RNA-pRNA product and the absence of the DNA-pDNA product revealed that the RNA but not the DNA construct is a substrate of zebrafish Rlig1 (Figure 1e), demonstrating selectivity of zebrafish Rlig1 for RNA. Moreover, we found that the ligation reaction is rapid, with 40% of product formation already achieved within 5 min of incubation (Figure 1e–f) congruent with previous work on human Rlig1^4^. In summary, zebrafish Rlig1 is able to ligate RNA hairpin substrates *in vitro* that possess a nick in their loop comprising a 3’-hydroxyl terminus and a 5’-phosphate end. Given that this RNA ligase activity is analogous to that of human Rlig1, we decided to use zebrafish as an experimentally accessible model to study the function and biological role of Rlig1 *in vivo*.

### Rlig1 mRNA is maternally contributed and localised in the brain of zebrafish larvae

In order to determine the expression pattern of *rlig1* throughout embryonic development, we assessed the relative levels of *rlig1* mRNA in several embryonic stages. Total RNA was extracted from a pool of embryos of the desired stage and subjected to a reverse transcription quantitative polymerase chain reaction (RT-qPCR). We designed TaqMan probes to bind *rlig1* or *elongation factor 1 α* (*eef1a*) mRNA, respectively, the latter of which served as a reference gene^15^. To obtain relative *rlig1* mRNA levels, we calculated the inverse ΔCq (1/(Cq*^rlig1^*-Cq*^eef1a^*)) for each developmental stage. Interestingly, *rlig1* mRNA levels were particularly high at the beginning of the cleavage period (2- to 64-cell stage), and decreased during the blastula period (sphere stage) until steadily low levels were reached at the shield stage (gastrula period) (Figure 2a). The relatively high *rlig1* mRNA levels during the cleavage period indicate maternal deposition of *rlig1* mRNA as the embryo lacks *de novo* transcription until the maternal-zygotic transition^16^, which typically occurs at the 512-cell stage for the majority of genes^17^. The onset of zygotic transcription is accompanied by the degradation of maternal mRNA^18^. The presence of *rlig1* mRNA at stages preceding the maternal-zygotic transition is therefore likely solely due to maternal deposition of *rlig1* mRNA into the egg.

**Figure 2.**
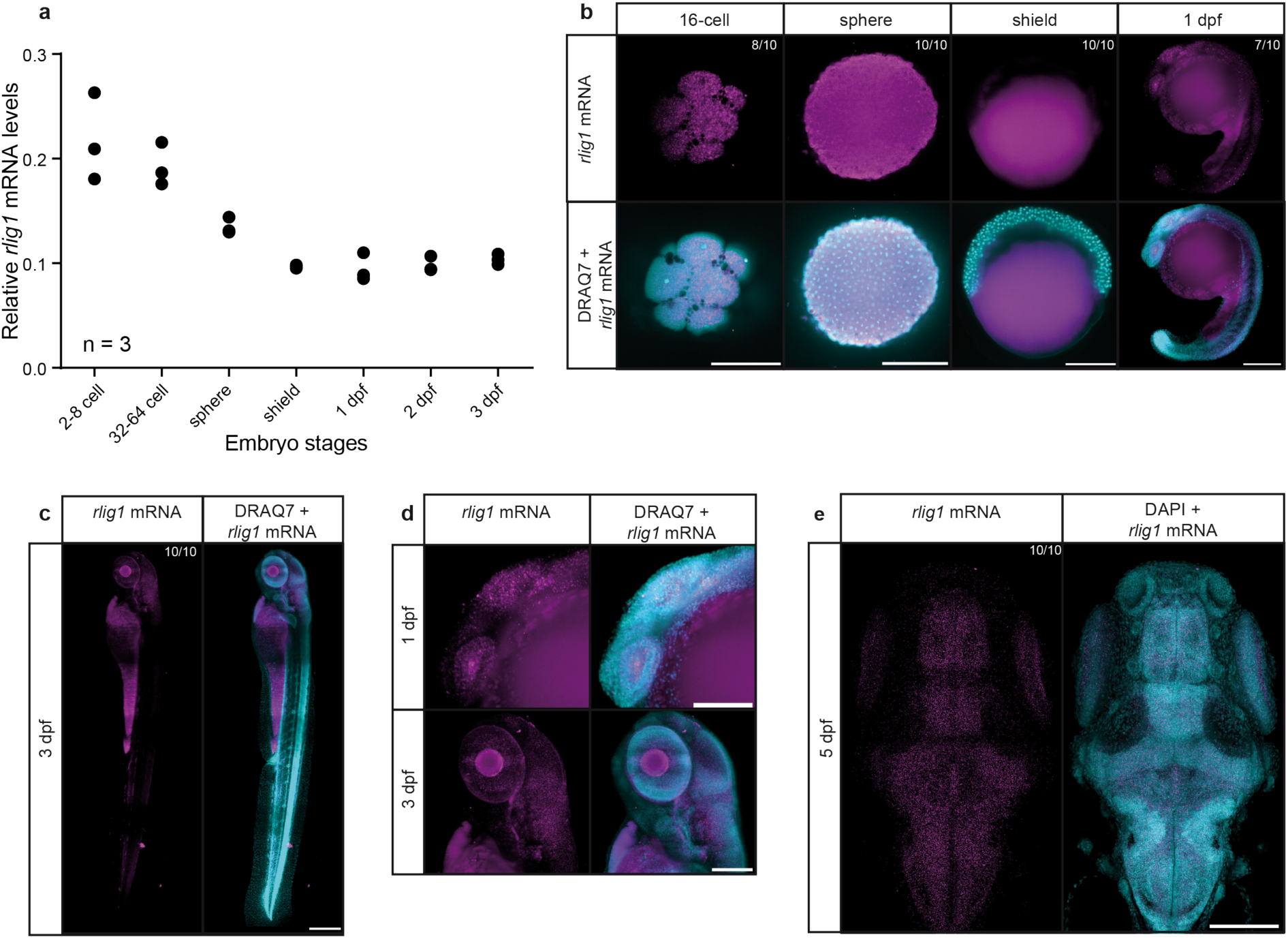
Expression analysis of *rlig1* mRNA during zebrafish embryonic development. **(a)** Relative *rlig1* mRNA levels at different developmental stages, measured by qPCR. The y-axis represents the inverse Cq value, and data are shown from triplicate experiments. **(b-d)** HCR RNA FISH images showing *rlig1* mRNA signal (magenta) and the nuclear reference signal (DRAQ7, cyan) at different developmental stages: (b) 16-cell stage (animal pole view, single optical slice), sphere stage (animal pole view, single optical slice), shield stage (lateral view, single optical slice), and 1 dpf (lateral view, anterior to the left, maximum intensity projection). Scale bar represents 250 µm. (c) 3 dpf embryo in full length (stitched image, maximum intensity projection). The embryo is shown in lateral view with anterior to the top. Scale bar represents 250 µm. (d) Enlarged views of the head region from embryos presented in (b) and (c) (maximum intensity projection). Scale bar: 150 µm. **(e)** *rlig1* (magenta) HCR RNA FISH image of a 5 dpf larva with DAPI (cyan) as a reference. The larva is shown in dorsal view with anterior to the top (maximum intensity projection). Scale bar: 150 µm. Representative image from one biological replicate (n = 1); additional samples imaged with an alternative microscopy technique showed the same expression pattern. The number of embryos showing a comparable expression pattern is indicated as n/10 in each panel (b), (c), and (e).

Next, we examined the spatial localisation of *rlig1* mRNA in embryos during development using Hybridisation Chain Reaction RNA Fluorescence In Situ Hybridisation (HCR RNA FISH)^19^. We fixed wild-type (WT) embryos at defined developmental stages and applied probe sets targeting *rlig1* together with DRAQ7 as a nuclear counterstain. To distinguish between auto-fluorescence or background signal and specific signal, we performed control experiments without *rlig1* probe (Supplementary Figure 1), which revealed auto-fluorescence of the yolk and parts of the eye (from 2 days post-fertilisation (dpf) onwards) manifesting as a blurred signal. In contrast, specific *rlig1* signals are clearly present in distinct intracellular puncta (Figure 2b–e) that are typical for mRNAs^20^.

For embryos of the cleavage and blastula period, *rlig1* transcripts were distributed evenly within the embryo (Figure 2b). In the gastrula period (shield stage), no *rlig1* mRNA signal could be detected in the embryo (Figure 2b), in agreement with our RT-qPCR data (Figure 2a). In the pharyngula period (1 dpf), *rlig1* transcripts were distributed throughout the embryo (Figure 2b) with the signal being strongest in the head, particularly in the eyes and brain (Figure 2b–d) and decreasing towards the tip of the tail. At the end of the hatching period (3 dpf), embryos displayed a notable accumulation of *rlig1* transcripts within the brain and eyes (Figure 2c–d), while only a minimal signal could be discerned in the tail region, parallel to the spinal cord (Figure 2c); similar observations were made for older larvae (5 dpf). In the brain, an even distribution of *rlig1* mRNA was observed, with no tendency for specific regions of the brain (Figure 2e). In addition, *rlig1* transcripts accumulated in the eyes. However, due to the auto-fluorescence of the eye, which increases with development (Supplementary Figure 1), this observation is difficult to quantify.

In summary, *rlig1* transcripts are evenly distributed throughout the embryo during early development (cleavage-blastula period), whereas they are not detectable above background in the shield-stage embryo (gastrula period). Starting at the pharyngula stage (1 dpf), an enriched localisation of *rlig1* transcripts can be observed, particularly in the brain and eyes. This raised the question as to whether *rlig1* is involved in the processing or transmission of visually perceived stimuli.

### CRISPR/Cas9-mediated rlig1 knockout alters behavioural response to visual stimuli

To gain insight into the biological role of Rlig1 *in vivo*, we generated zebrafish *rlig1* mutants using a CRISPR/Cas9-mediated strategy to remove the promoter region, the first exon and part of the second exon of the *rlig1* gene (*zgc:103499*) (Figure 3a). The successful *rlig1* knockout was verified on the genomic level by PCR spanning the deletion (Supplementary Figure 2a) and on the transcript level by mRNA sequencing, RT-qPCR (Figure 3b), and HCR RNA FISH (Figure 3c). As intended, the deletion of the extended *rlig1* promoter-region resulted in a *rlig1* null mutant (Supplementary Figure 2b) not producing any *rlig1* mRNA (Figure 3b–c).

**Figure 3.**
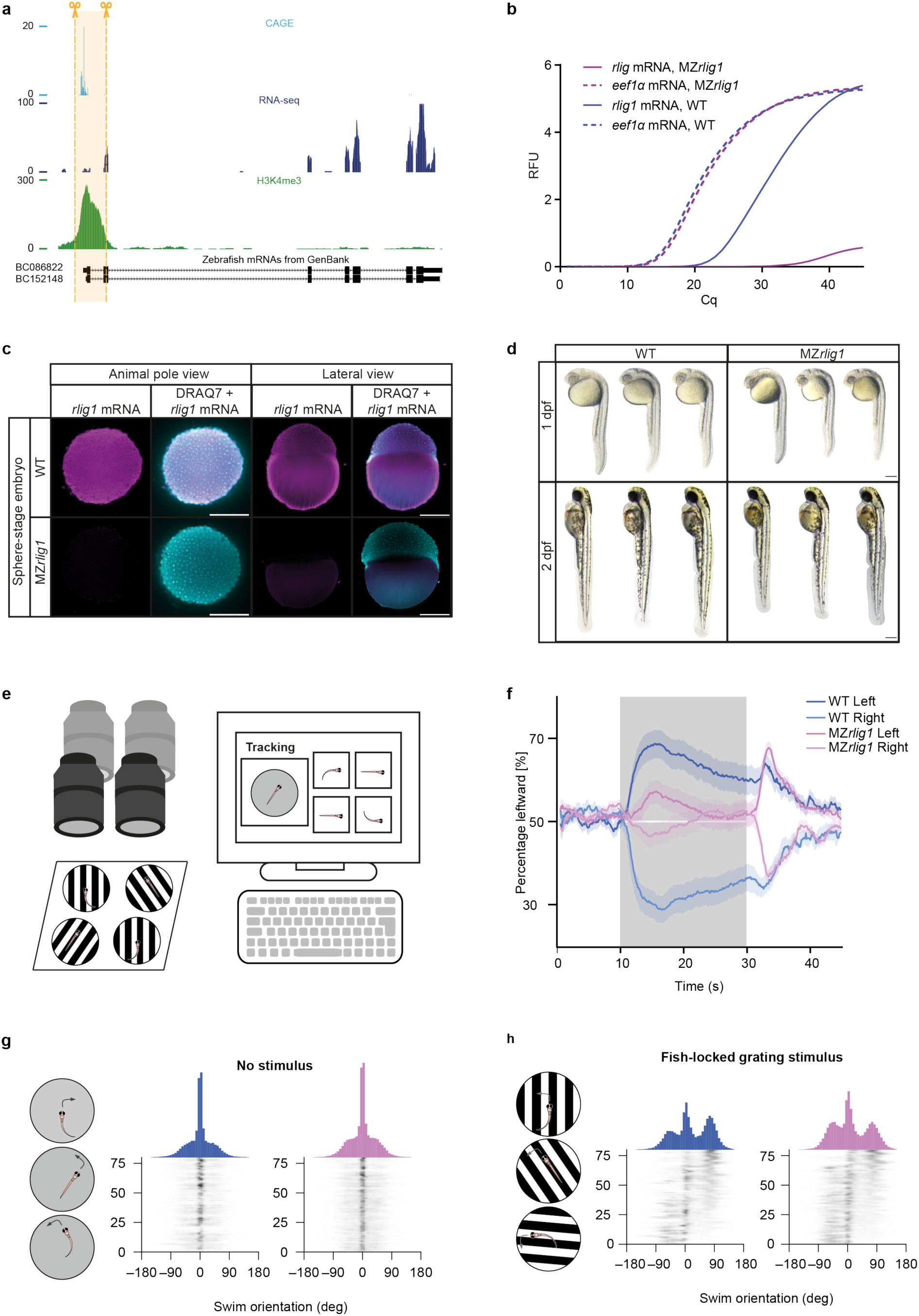
Generation and validation of *rlig1^-/-^* mutants and behavioural analysis of maternal-zygotic *rlig1^-/-^* larvae. **(a)** Generation of a CRISPR/Cas9-mediated *rlig1* mutant. Schematic of the *rlig1* locus, including associated RNA-Seq, H3K4me3, and CAGE data at the dome to 30% epiboly stage^56^. CAGE data (light blue) indicate observed transcription start sites of mRNAs, while H3K4me3 (green) represents a histone modification enriched at active promoters. RNA-Seq data (dark blue) provide information on transcribed sequences. The *rlig1* mRNA (black) connects the displayed data to coding (exonic) and non-coding (intronic) regions. The shaded orange area marks the genomic region targeted for deletion by CRISPR/Cas9-based mutagenesis. **(b-c)** Validation of the gene knockout in 5 dpf maternal-zygotic *rlig1^-/-^* mutant larvae. (b) RT-qPCR analysis with relative fluorescence units (RFU) on the y-axis and Cq-values on the x-axis. Representative plot of n = 3. Total RNA was extracted from a pool of ten 5 dpf larvae. (c) HCR RNA FISH analysis of WT and MZ*rlig1* embryos. Representative animal pole (left) and lateral (right) views of WT (upper row) and MZ*rlig1* (lower row) embryos at the sphere stage show the *rlig1* mRNA (magenta) and the nuclear reference signal (DRAQ7, cyan) (WT n = 10, MZ*rlig1* n = 5). Scale bar: 250 µm. **(d)** Morphology of WT and MZ*rlig1* embryos at 1 dpf and 2 dpf. Shown are lateral views with anterior to the left. Scale bar represents 250 μm. **(e)** Schematic representation of the behavioural experiment setup. A zebrafish larva (brown) is positioned above a projected visual stimulus. A camera mounted above the larva tracks its movement within the experimental arena, allowing real-time data processing and storage. This setup enables closed-loop experiments, in which the projected stimulus adapts dynamically to the larva’s position. **(f)** Responsiveness of larvae to leftward or rightward motion stimuli. The percentage of leftward swimming responses (y-axis) is plotted over time (x-axis) for WT (blue) and MZ*rlig1* (pink) larvae. Dark blue and dark pink lines represent responses to a leftward stimulus, while light blue and light pink lines correspond to responses to a rightward stimulus. The stimulus presentation period (10–30 s) is indicated by a grey-shaded background. The average response across all larvae is shown as a solid line, with the shaded area representing the standard error of the mean (SEM). n = 80 for MZ*rlig1* and WT larvae. **(g,h)** Swim orientation of WT and MZ*rlig1* larvae in the absence or presence of a directional stimulus. Swim orientation (in degrees) of individual bouts is shown for WT (blue) and MZ*rlig1* (pink) larvae. (g) Swim orientation in the absence of a stimulus. (h) Swim orientation during presentation of a leftward-motion stimulus. n = 80 for MZ*rlig1* and WT larvae.

Neither zygotic nor maternal-zygotic *rlig1*^-/-^ mutants (MZ*rlig1* mutants) showed obvious development defects, although a slight reduction in overall body length was observed in MZ*rlig1* embryos (Figure 3d). Since *rlig1* mRNA is highly abundant in the larval brain and eyes, we decided to assess the behaviour of MZ*rlig1* larvae. We chose the optomotor response as it is a complex behaviour requiring the spatial and temporal integration of sensory input, computation of movement direction, and control of motor output, making it a sensitive assay for detecting perturbations in neuronal function. To this end, we presented the larvae with a moving grating stimulus and tracked their movement in response to it (Figure 3e). If no stimulus is presented, larvae display random swim bouts to the left and right as well as forward. Upon presentation of a left- or rightward motion stimulus, relative to the body axis of the animal, it has been shown that larvae will follow the direction of the stimulus^21,22^. Analysis of the response, averaged over all repetitions during the stimulus presentation of each larva and over all individuals, showed that in approximately 70% of trials, WT larvae followed the stimulus, for both tested directions (Figure 3f). In contrast, in only around 55% of trials MZ*rlig1* larvae followed the presented stimulus (Figure 3f). The mean random swimming direction of the larvae when no stimulus was presented fluctuated around chance levels of 50% (Figure 3f). Therefore, a response of 55% is evidence for a strongly reduced behaviour. This can also be deduced from the analysis of the following tendencies over the time course of stimulus presentation. In the absence of a stimulus, most swim bouts are directed forward for both WT and MZ*rlig1* larvae (Figure 3g). Presentation of a grating stimulus, like in the depicted example with leftward motion, is expected to decrease the total number of forward bouts and increase the frequency of bouts following the stimulus. This behaviour was indeed observed in WT larvae, where the peak of leftward bouts was nearly twice as high as for rightward bouts (3181 vs 1851 bouts, Figure 3h, left panel). For MZ*rlig1* larvae, the number of forward bouts decreased upon stimulus presentation, but the peak of leftward bouts did not exceed the peak of rightward bouts (2663 vs 2756 bouts, Figure 3h, right panel). Thus, upon stimulus presentation MZ*rlig1* larvae perform fewer bouts following the stimulus direction (37% left swims) than WT (45% left swims). Interestingly, the behavioural response following the direction of the stimulus increased shortly after the end of the stimulation period for both WT and MZ*rlig1* larvae (Figure 3f). This suggests that the reduced response of MZ*rlig1* larvae to the grating stimulus is unlikely to result solely from a general impairment of the visual system, as they exhibit a delayed but stimulus-contingent behavioural response following stimulus offset. This observation indicates that basic visual perception is at least partially preserved. Moreover, normal baseline swimming behaviour of the MZ*rlig1* larvae supports the assumption that motor output is likely intact (Supplementary Figure 3). This suggests that the underlying deficit may originate downstream of primary sensory detection, potentially at the level of signal processing or sensory-motor integration.

### Absence of Rlig1 alters neuronal activity in the anterior hindbrain

To investigate whether the deficit in behavioural response originates at the level of visual signal processing, sensory–motor integration, or motor output, we examined neuronal activity in three brain regions known to be involved in the transmission and processing of visual stimuli in larval zebrafish: the pretectum, tectum, and anterior hindbrain (aHb)^23,24^. Perceived visual information travels from the retina through the retinal ganglion cells to one of 10 retinal arborization fields (AFs), depending on the nature of the presented visual stimulus^25^. Motion and direction-related information is sent to AF5 (pretectum) and AF10 (tectum), with the tectum receiving input from the majority of retinal ganglion cells^26,27^. We focused on the signal transmission route from the pretectum to the anterior hindbrain, where motion-related information is integrated and relayed to motor output neurons^23,28,29^. It is important to note that visual input can also reach the aHb via the tectum, which serves as an alternative processing centre for motion information^26,27^. To account for this, we additionally examined neuronal activity in the tectum (midbrain) as a reference, allowing us to assess whether signals might be transmitted through this parallel route.

To determine whether the altered MZ*rlig1* behaviour can be explained by a sensory, motor, or sensory-motor transformation deficit, we imaged the neural activity upon motion stimulus presentation (Figure 4a). For this, we used a setup similar to the optomotor response assay (Figure 3e), except that the larvae were immobilized and exposed to left- and rightward moving dot stimuli, while neural activity of all neurons within a selected imaging plane was recorded simultaneously. One plane each in the aHb and in the pretectum was selected for Ca^2+^-imaging of *Tg(elavl3:H2B-GCaMP8s)^mpn43830^*;*mitfa^-/-^*;*rlig1*^-/-^ and *Tg(elavl3:H2B-GCaMP8s) ^mpn438^;mitfa^-/-^* larvae. Nuclear-localised GCaMP8s expression levels across the brain were comparable between genotypes. Interestingly, neural activity across trials in direction-selective regions of the pretectum and the aHb was reduced in MZ*rlig1* larvae (Figure 4b–d). Although both regions showed a significant reduction, the effect size was considerably larger in the aHb compared to the pretectum (Cohen’s d −0.742 vs −0.398). The temporal dynamics of the MZ*rlig1* pretectal and aHb activity qualitatively matched the WT with fast dynamics in the pretectum and slow integration times in the aHb^28^ (Figure 4c–d).

**Figure 4.**
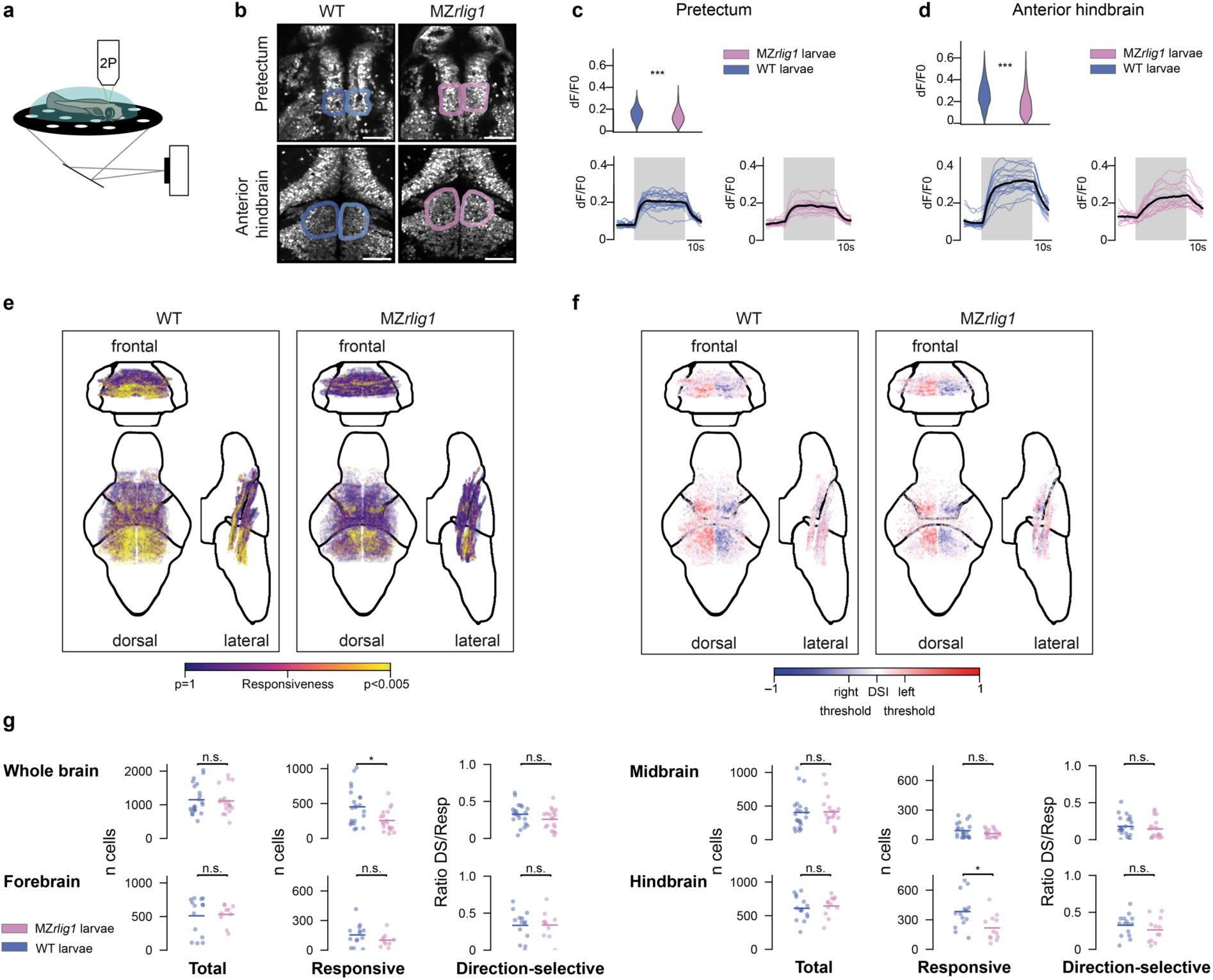
Imaging of neuronal activity in the forebrain, midbrain, and hindbrain of WT and *MZrlig1* larvae using two-photon imaging. **(a)** Schematic representation of the experimental setup. A motion stimulus is projected onto the surface under a larva fixed in agarose while a selected plane of a specific brain region is imaged simultaneously. **(b)** The neurons in the pretectum (upper row) and anterior hindbrain (lower row) are shown, with the motion-responsive regions considered for analysis marked in blue (WT) and pink (MZ*rlig1*). Scale bar: 50 µm **(c)** Representation of neural activity across trials of motion-responsive regions in the pretectum (top). WT larvae are shown in blue (n = 10) and MZ*rlig1* (n = 9) larvae in pink. Neuronal activity is represented as dF/F0. For statistical analysis, a one-sided t-test was performed (t = –5.85, p = 3.481 × 10⁻^9^, Cohen’s d = –0.398), and significance is indicated: *** corresponds to p < 0.001. The temporal dynamics of pretectal activity are shown in the lower panel. Blue lines represent the neural activity of WT larvae across trials, with the black line representing the mean. The same representation is shown in pink for MZ*rlig1* larvae. The shaded grey area indicates the period during which the stimulus was presented to the larvae. **(d)** Representation of neural activity across trials of motion-responsive regions in the anterior hindbrain (top). WT larvae are shown in blue (n = 9) and MZ*rlig1* larvae in pink (n = 8). Neuronal activity is represented as dF/F0. For statistical analysis a one-sided t-test was performed (t = –10.55, p = 8.993 × 10⁻^25^, Cohen’s d = –0.742), and significance is indicated: *** corresponds to p < 0.001. The temporal dynamics of anterior hindbrain activity are shown in the lower panel. Blue lines represent the neural activity of WT larvae across trials, with the black line representing the mean. The same representation is shown in pink for MZ*rlig1* larvae. The shaded grey area indicates the period during which the larvae were presented with the stimulus. **(e)** Depiction of each neuron in WT (left) and MZ*rlig1* (right) larvae across the imaged brain regions. The colour of the dots corresponds to the neuron’s responsiveness to the stimulus, with purple representing no motion-responsiveness and yellow indicating high motion-responsiveness (p < 0.005). The brain is depicted in frontal (top), dorsal (left), and lateral (right) views to show the distribution and depth of imaged neurons. **(f)** Representation of motion-responsive neurons in WT (left) and MZ*rlig1* (right) larval brains according to their direction-selectivity. The brain is depicted in frontal (top), dorsal (left), and lateral (right) views to show the distribution and depth of imaged neurons. Direction-selective neurons are highlighted in colour with neurons responsive to rightward stimuli marked in blue, and neurons responsive to leftward stimuli marked in red. DSI: Direction selectivity index. **(g)** Representation of the total number of neurons (left), the number of motion-responsive neurons (middle), and the ratio of direction-selective (DS) to all motion-responsive neurons (right). WT larvae are shown in blue and MZ*rlig1* larvae in pink. Dots represent the number of neurons or the ratio of direction-selective to motion-responsive neurons in individual larvae, with the blue or pink lines representing the mean. For statistical analysis, a one-sided t-test was performed with Bonferroni correction, and a significant difference was determined for motion-responsive neurons in the whole brain (t = –2.865, p = 0.003, Cohen’s d = –0.897) and specifically in the hindbrain (t = – 2.772, p = 0.005, Cohen’s d = –1.073); * = p < 0.05. Sample sizes: MZ*rlig1* hindbrain: 12 planes from 11 larvae, MZ*rlig1* midbrain: 18 planes from 17 larvae, MZ*rlig1* forebrain: 10 planes from 10 larvae; WT hindbrain: 14 planes from 12 larvae, WT midbrain: 22 planes from 20 larvae, and WT forebrain: 13 planes from 11 larvae.

Next, we compared the total number of neurons, number of motion-responsive neurons, and number of direction-selective neurons in WT and MZ*rlig1* larvae. No significant difference between the total number of automatically segmented neurons^31^ in WT and MZ*rlig1* larvae across the imaged brain regions was observed (Figure 4e–g). However, a more detailed analysis of the number of responsive neurons revealed that MZ*rlig1* larvae exhibited a reduced number of motion-responsive hindbrain neurons in comparison to their WT relatives (Figure 4e–g). Nevertheless, the fraction of direction-selective neurons among these motion-responsive neurons was found to be unaltered in comparison to their WT controls (Figure 4f–g). Furthermore, the direction-selective neurons in WT larvae did not have higher neural activity than the direction-selective neurons in MZ*rlig1* larvae (Supplementary Figure 4). This suggests that the reduced neural activity across the aHb is a result of a reduced number of motion-responsive neurons. We did not find any differences in motion-responsiveness or direction selectivity in the forebrain (pretectum) and midbrain (tectum) (Figure 4g).

These data suggest that the observed difference in the number of motion-responsive neurons in the aHb is due to a deficit in the transmission of the signal between the pretectum and the aHb causing fewer neurons in the aHb to be activated upon stimulus presentation. It may therefore be presumed that the observed behavioural response is not solely attributable to a sensory-perception deficit or a motor output deficit (Supplementary Figure 3), but rather due to a processing-transmission deficit. It is possible that this involves alterations in the molecular profile of neurons that disrupt their communication dynamics, or impairments in other cellular mechanisms across the body. To test this, we analysed the impact of *rlig1* loss on global transcriptional profiles across the whole animal.

### rlig1 mutants have drastically altered developmental transcriptomes

To understand the neurodevelopmental origins of the observed behavioural and neuronal deficits at the gene regulatory level, we performed transcriptome profiling of MZ*rlig1* embryos at multiple developmental stages and compared their gene expression patterns to WT controls. The goal was to capture dynamic gene expression changes throughout embryogenesis and identify downstream pathways potentially affected by the absence of Rlig1. We selected representative developmental stages – 4-cell stage, sphere stage, shield stage, bud stage, 1 dpf, and 5 dpf – and extracted total RNA from pools of 8-10 age- and genotype-matched embryos for poly(A) enrichment, sequencing, and differential gene expression analysis. The *rlig1* expression dynamics captured by RNA-Seq across these developmental stages closely matched the spatiotemporal patterns previously observed by qRT-PCR and HCR RNA FISH (Supplementary Table 1).

Principal component analysis (PCA) revealed clear clustering along PC1 according to developmental time, ranging from the 4-cell stage to 5 dpf (Figure 5a), confirming that embryonic age was the main driver of transcriptomic variance. This developmental progression was preserved in both WT and MZ*rlig1* samples. Interestingly, differential gene expression analysis revealed extensive transcriptional changes in *rlig1*-deficient embryos across developmental stages, with the highest number of differentially expressed genes (DEGs) detected at the sphere stage (>500 DEGs; Figure 5b). Although some genes were differentially expressed across multiple time points (Figure 5c–d), most DEGs were stage-specific (Figure 5e). The number of upregulated genes generally exceeded the number of downregulated genes (|log₂FoldChange| ≥ 1.5, padj ≤ 0.05) (Figure 5b). The lack of overt morphological phenotypes during embryogenesis (Figure 3d), despite the widespread transcriptional alterations in *rlig1*-deficient embryos, could indicate functional redundancy through a paralog. However, as no Rlig1 paralog could be identified in the zebrafish genome based on amino acid sequence homology, the compensatory mechanisms are likely unrelated to functional redundancy. Instead, the pronounced transcriptional changes may reflect the activation of alternative compensatory pathways.

**Figure 5.**
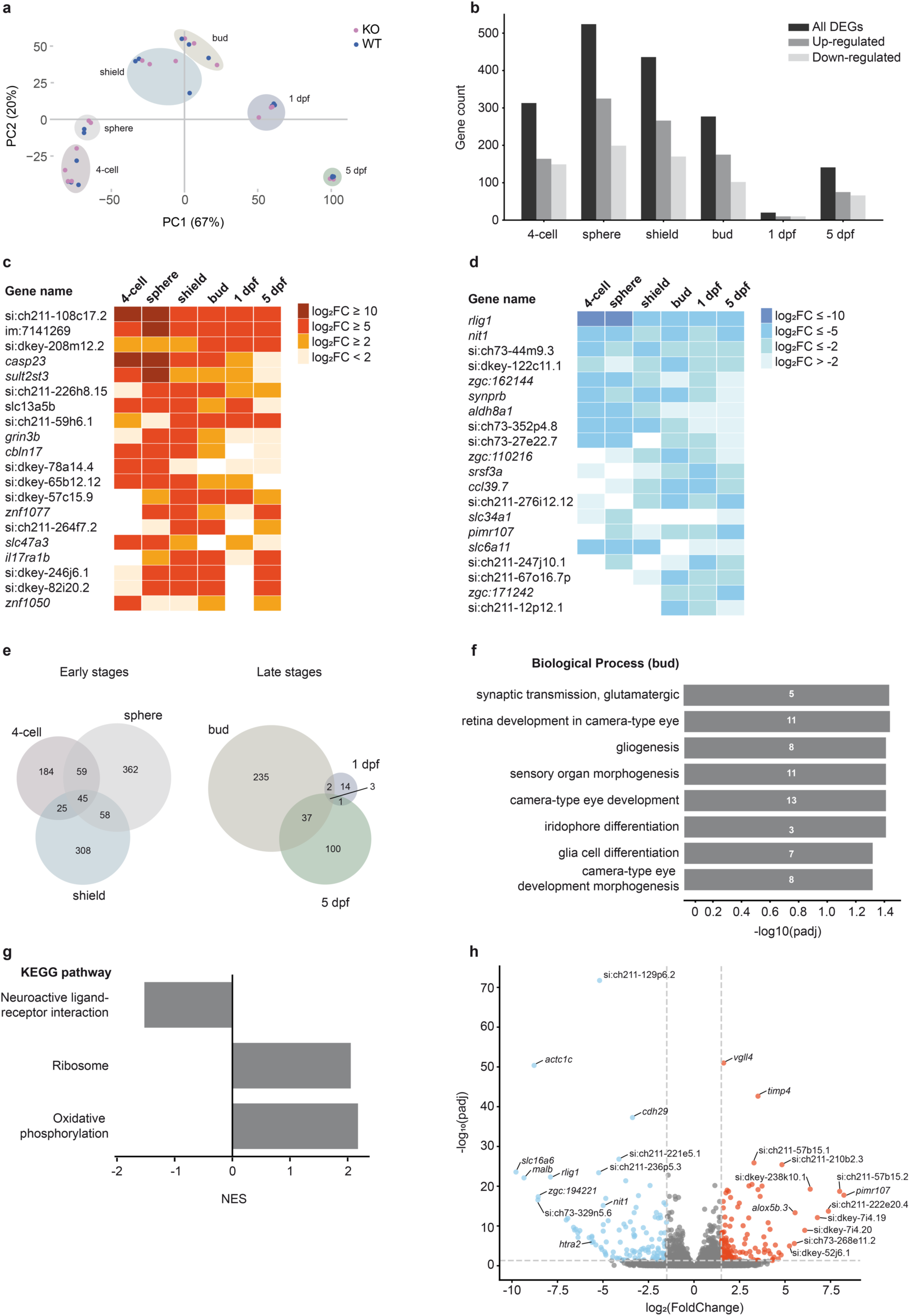
Transcriptomic profiling reveals stage-specific gene expression changes in *rlig1*-deficient embryos. **(a)** PCA of RNA-Seq samples across all developmental stages. Each dot represents one biological replicate. WT samples are shown in blue, MZ*rlig1* (KO) samples in pink. Samples of the same developmental stage are indicated by shaded background areas. **(b)** Number of differentially expressed genes (DEGs) per developmental stage (adjusted p-value ≤ 0.05, |log₂FC| ≥ 1.5). Upregulated genes in MZ*rlig1* are shown in grey, downregulated genes in light grey, all DEGs combined in dark grey. **(c)** Matrix of the 20 most frequently upregulated genes across stages. Genes were included if they appeared among the top 50 upregulated DEGs (based on log₂FC) in at least two stages. Colour intensity reflects the log₂FC value (≥ 0), from light orange to dark red. White cells indicate that the gene was not among the top 50 upregulated genes in that stage. **(d)** Matrix of the 20 most frequently downregulated genes, corresponding to the criteria in (c). Colour intensity reflects the log₂FC value (≤ 0), from light blue to dark blue. White cells indicate absence from the top 50 downregulated genes in the respective stage. **(e)** Venn diagrams showing the overlap of DEGs between adjacent developmental stages. Up-and downregulated genes are combined. The left panel displays early stages (4-cell, sphere, shield), the right panel later stages (bud, 1 dpf, 5 dpf). **(f)** GO enrichment analysis of DEGs at the bud stage (category: biological process). The x-axis shows the –log₁₀-transformed adjusted p-value; the y-axis lists significantly enriched terms. Numbers shown on the bars represent the number of DEGs associated with each term. **(g)** Gene Set Enrichment Analysis (GSEA) of combined-stage transcriptomes, comparing MZ*rlig1* and WT embryos. Shown are significantly enriched KEGG pathways (adjusted p < 0.05), ranked by normalised enrichment score (NES). **(h)** Volcano plot showing differentially expressed genes in dissected heads of 5 dpf MZ*rlig1* compared to WT larvae. Each point represents a gene, plotted by log₂(FoldChange) and – log₁₀(adjusted p-value). Genes meeting the significance threshold (|log₂FC| ≥ 1.5, padj ≤ 0.05) are coloured: significantly upregulated genes in red, downregulated genes in blue, and non-significant genes in grey. Selected genes are labelled.

The identified DEGs functionally grouped into several categories, including RNA processing, immune response, intracellular and transmembrane transport, and synaptic signalling (Supplementary Tables 2–7). Strikingly, a notable number of solute carrier family genes were differentially expressed, including *slc13a5b* and *slc47a3*, which were consistently upregulated across several stages (Figure 5c). Genes involved in neurotransmitter transport and synaptic signalling, such as *grin3b*^32,33^ and *slc6a11*^34^ were also affected – even at very early stages of neurodevelopment (Figure 5c sphere-bud and Figure 5d 4-cell-shield, respectively). A detailed gene ontology (GO) analysis revealed distinct, stage-specific enrichments of biological processes in *rlig1*-deficient embryos (Supplementary Tables 2–7), indicating that Rlig1 influences a broad spectrum of cellular functions during development. Ranging from mitochondrial electron transport and redox activity at the 4-cell and sphere stages, to immune-related and cell cycle-associated processes at 1 dpf, and transmembrane transport functions across several stages, the affected pathways reflect a dynamic and temporally coordinated transcriptomic response to Rlig1 loss.

In particular, processes associated with synaptic signalling and brain development emerged across multiple stages, with the bud stage exhibiting the most pronounced transcriptional signature. Differentially enriched GO terms included “synaptic transmission, glutamatergic”, “gliogenesis”, “glial cell differentiation”, and “sensory organ morphogenesis” (Figure 5f), supporting the hypothesis that Rlig1 may contribute to early neurodevelopmental processes. Notably, we also observed enrichment of several GO terms related to eye development – such as “retina development in camera-type eye” and “camera-type eye morphogenesis” – during the bud stage (Figure 5f, Supplementary Table 5). This aligns with the onset of eye-field specification, which begins around 10 hpf and corresponds to early molecular patterning events in the developing visual system^35,36^. While our behavioural results point to impaired neural processing as the primary cause of the sensorimotor deficits, these transcriptomic findings suggest that early aspects of visual system development may also be affected. In particular, potential alterations in retinal ganglion cells, which transmit visual information from the retina to the arborization fields in the brain^37,38^, could contribute to the observed phenotype.

To move beyond stage-specific effects and assess broader, genotype-dependent changes, we performed Gene Set Enrichment Analysis (GSEA) using Kyoto Encyclopedia of Genes and Genomes (KEGG) pathway annotations (Supplementary Tables 2–7). This analysis, which aggregated transcriptome data across all developmental stages, revealed three pathways that were consistently and significantly enriched in MZ*rlig1* mutants compared to WT embryos: “oxidative phosphorylation”, “ribosome”, and “neuroactive ligand-receptor interaction” (Figure 5g). Notably, components of neuroactive ligand-receptor interactions were predominantly downregulated, whereas genes associated with oxidative phosphorylation and ribosome pathways were largely upregulated. These results suggest that Rlig1 deficiency exerts stable, stage-independent effects on key biological processes, including mitochondrial function, protein biosynthesis, and neurotransmitter signalling, which may underlie the neural and behavioural deficits we observed.

To better understand the transcriptional basis of the altered behavioural phenotype observed in *rlig1*-deficient larvae, we performed RNA sequencing on dissected heads of 5 dpf larvae, where we found *rlig1* expression to be enriched (Figure 2e). We classified larvae as either “good” or “bad” responders based on the number and directionality of swim bouts in our optomotor stimulus paradigm (Figure 3g). By combining behavioural categorisation with transcriptome profiling, we aimed to determine whether stimulus responsiveness is associated with distinct gene expression patterns. DEG analysis revealed substantial transcriptional changes between genotypes (Figure 5h), but no consistent correlation with behavioural responsiveness (Supplementary Figure 5, Supplementary Table 8). Nevertheless, the head transcriptomes of MZ*rlig1* larvae contained numerous differentially expressed genes that are associated with categories also altered in the developmental dataset. Among the significantly downregulated genes identified in 5 dpf larval heads, *slc16a6* exhibited the strongest reduction in transcript levels (log₂FoldChange (FC) = –9.78) (Figure 5h). As a member of the solute carrier family, it exemplifies a broader pattern of disrupted transmembrane transport processes also observed in the developmental dataset. Another gene of interest is *htra2*, which encodes HtrA serine peptidase 2, a mitochondrial protease involved in maintaining protein homeostasis^39^ (Figure 5h). Loss of HtrA2 function has been linked to neurodegenerative disease pathways, particularly Parkinson’s disease, and triggers a brain-specific transcriptional stress response^40^. These findings reinforce the conclusion that loss of Rlig1 impacts both solute carrier–mediated transport and neuronal signalling pathways, consistent with the developmental transcriptome data. While our RNA-Seq data revealed genome-wide expression changes and identified consistent enrichment of specific pathways across stages, future studies will be valuable for validating the expression patterns of individual genes.

## Discussion

In this study, we examined the biological function of zebrafish Rlig1, an RNA ligase that catalyses the ligation of nicked RNA hairpin substrates *in vitro*. We found that *rlig1* transcripts are maternally deposited and broadly distributed throughout the early embryo, with highest levels during the cleavage period. Transcript abundance declined toward gastrulation, and from 1 dpf onwards, *rlig1* mRNA exhibited enriched localisation in the brain and eyes.

To investigate the *in vivo* relevance of this expression pattern, we generated *rlig1* knockout zebrafish. Despite the absence of overt morphological abnormalities during embryogenesis, and the lack of detectable phenotypes in acoustic startle response behaviour in a previous study^41^, MZ*rlig1* larvae exhibited a reduced behavioural response to visual motion stimuli. This phenotype was accompanied by diminished stimulus-evoked neuronal activity in both the pretectum and the anterior hindbrain. To gain insight into the molecular mechanisms underlying these phenotypes, we profiled the transcriptomes of WT and MZ*rlig1* embryos across six developmental stages. The data revealed widespread gene expression changes in mutants, many of which were stage-specific, while others were shared across time points. GO analyses further indicated temporally distinct enrichment of neurodevelopmental pathways. First, differential expression of several components of the FGF signalling cascade – including *dusp* and *fgfr* genes (Supplementary Tables 9–14) – suggests that Rlig1 loss affects pathways involved in midbrain-hindbrain patterning^42,43^, regions in which we observed functional deficits. Supporting this idea, a study in human cells demonstrated that Rlig1 can interact with phosphorylated ERK to activate mTORC1 signalling^44^, linking Rlig1 to broader growth-regulatory and neurodevelopmental networks, consistent with the role of FGF receptors as upstream activators of the MAPK/ERK pathway^45^. Second, among the most significantly enriched terms were “synaptic transmission” and “retina development in the camera-type eye”. These enrichments point toward early disturbances in brain and visual system development. While impaired neural processing is a likely contributor to the observed behavioural deficits, the transcriptomic findings raise the possibility that upstream alterations – such as in retinal ganglion cell development – may also play a role. Taken together, the concordant effects across behavioural assays, brain-wide calcium imaging, and transcriptomic profiling argue for a direct link between loss of Rlig1 and disrupted visual processing.

Strikingly, our pathway analysis across all developmental stages revealed consistent enrichment of oxidative phosphorylation, ribosome function, and neuroactive ligand-receptor interaction in *rlig1*-deficient embryos. These findings reflect stable, genotype-dependent effects on mitochondrial function, translation, and neurotransmitter signalling. Among these, the persistent enrichment of the ribosome pathway is particularly notable, as it points to a potential role of Rlig1 in translational control. This interpretation is consistent with previous findings in human cells, where knockout of Rlig1 led to reduced ribosome integrity and increased RNA degradation under oxidative stress^4^. Moreover, recent biochemical studies have identified Rlig1 as an atypical RNA ligase capable of ligating tRNAs *in vitro*, particularly within the anticodon loop, and binding tRNA *in vivo*^11^. In support of this, transcriptomic analyses of Rlig1-deficient mouse brains revealed altered global tRNA levels, further implicating Rlig1 in tRNA metabolism^11^. Although our poly(A)-enriched RNA-Seq data do not capture tRNAs or rRNAs, the observed upregulation of translation-related pathways in *rlig1* mutants may reflect compensatory responses to translational stress or disrupted RNA homeostasis. Neurons in active processing circuits, such as those for vision, have exceptionally high metabolic and translational demands^46,47^, which may render them exquisitely sensitive to disruptions in core pathways like oxidative phosphorylation and ribosome function.

Many neurons, such as retinal ganglion cells, extend axons over long distances that can span several centimetres to more than a metre^48^. Such extreme cellular geometry requires local mRNA translation within axons to dynamically tailor the local proteome in response to changing physiological states or extracellular signals^48–50^. RNA repair activities mediated by RNA ligases could be particularly advantageous, as they would allow damaged transcripts to be maintained locally rather than relying on *de novo* transcription and long-distance transport from the soma. Without such repair, defective mRNAs in distal axons could impair local protein synthesis, compromise neuronal function, and contribute to neurological disorders^49^. The high mitochondrial content of neurons renders them a major source of reactive oxygen species^46,47^, which can oxidatively damage RNA, a process implicated in the pathogenesis of neurodegenerative disorders such as Alzheimer’s and Parkinson’s disease^51^. Therefore, our findings raise the intriguing possibility that Rlig1 contributes to neuronal resilience by preserving the integrity of axonally localised mRNAs under oxidative stress, thereby supporting mitochondrial function, synaptic signalling, and overall circuit stability. Consistent with this interpretation, the Comparative Toxicogenomics Database (CTD) associates the human Rlig1 orthologue with several neurological and cognitive conditions, including neurodevelopmental and neuroinflammatory disorders, Alzheimer’s disease, and cognition disorders, based on curated and inferred chemical–gene–disease relationships^52^.

Taken together, our findings establish Rlig1 as a previously unrecognised regulator of vertebrate development, acting at the intersection of RNA metabolism, signalling cascades, and neural circuit formation. While our transcriptomic analyses provide an integrated view of the regulatory changes associated with Rlig1 deficiency, the precise molecular substrates and interactors of Rlig1 remain to be identified. Given the use of bulk tissue and poly(A)-enriched RNA libraries, non-polyadenylated RNA species and cell-type-specific effects may have been missed. Future work using single-cell transcriptomics, RNA interactome mapping, or non-poly(A) capture techniques will be crucial to dissect the molecular functions of Rlig1 *in vivo*. Our study provides a foundation for understanding how RNA ligation contributes to the robustness of vertebrate neurodevelopment and function.

## Data and code availability

The RNA-Seq data has been deposited at the GEO repository (accession number: GSE308510) and can be accessed at https://www.ncbi.nlm.nih.gov/geo/query/acc.cgi?acc=GSE308510. The raw images, data, and source code for custom scripts used in this work are available from the corresponding authors upon request.

## Acknowledgments

This project has received funding from the European Research Council (ERC) under the European Union’s Horizon 2020 research and innovation program (ERC Advanced Grant #101019280 – AMP-Alarm to A.M., ERC Starting Grant #101075541 – CollectiveDecisions to A.B., and ERC Consolidator Grant #863952 – ACE-OF-SPACE to P.M.), the Emmy Noether Program (BA 5923/1-1 to A.B.), as well as the Deutsche Forschungsgemeinschaft (DFG, German Research Foundation) under Germany’s Excellence Strategy (EXC 2117 – 422037984). The Zukunftskolleg Konstanz provided additional project funds. K.S. was further supported by a Boehringer Ingelheim Fonds PhD fellowship and bridge funding provided by the International Max Planck Research School for Quantitative Behaviour, Ecology and Evolution (IMPRS-QBEE). F.S.K. thanks the Konstanz Research School Chemical Biology for support. We acknowledge the use of imaging equipment and expert support in microscope usage provided by Martin Stöckl and the Bioimaging Center at the University of Konstanz, as well as the technical help with setup building by our central university workshop. We also thank the animal welfare team at the University of Konstanz for their valuable support, and Andreas Gruber and Ian Kouzel for insightful discussions and expert advice on the analysis of the transcriptome data.

## Author contributions

Conceptualization: FSK, ACK, KS, AM, AB, PM; data curation: FSK, KS, HN; formal analysis: FSK, KS, OÖ; funding acquisition: AM, AB, PM; investigation: FSK, KS; supervision: ACK, AM, AB, PM; visualization: FSK, KS; writing – original draft: FSK, ACK, KS, AB, PM.

## Materials and Methods

### Expression and purification of recombinant zebrafish Rlig1

To obtain the zebrafish Rlig1 protein for functional ligase activity assay, an optimised sequence coding for zebrafish Rlig1 fused to an N-terminal His_6_-tag was ordered from GENEWIZ. Subsequent ligation, catalysed by a T4 ligase, introduced this coding sequence into a pET21b expression vector. The resulting pET21b-Rlig1-plasmid was transformed into *E. coli* BL21 RIL (DE3) competent cells. Expression and purification of the recombinant Rlig1 followed the protocol established by Yuan *et al.*^4^ for human Rlig1 with only minor adjustments. First, the transformed cells were cultured overnight in 50 mL LB medium containing 100 μg/mL carbenicillin at 37°C, 180 rpm. The next morning, the overnight culture was used to inoculate 1 L of LB medium containing 100 μg/mL carbenicillin to OD600 = 0.1. The suspension was incubated at 37°C at 180 rpm until it reached OD600 = 0.7. This was followed by cooling down the mixture on ice and subsequent induction of gene expression with 1.0 mM IPTG at 20°C for 18 h at 180 rpm. Cells were harvested by centrifugation at 8000 × g for 30 min at 4°C. The pellet was resuspended in 20 mL ice-cold lysis buffer (50 mM Tris-HCl pH 7.8, 500 mM NaCl, 2.0 mM β-mercaptoethanol, 10% (v/v) glycerol, 10 mM imidazole, cOmplete EDTA-free Protease inhibitor cocktail tablet) and lysed by sonication on ice. Next, the lysate was centrifuged at 40,000 × g for 30 min at 4°C and subsequently passed through a 0.45 μm syringe filter. The first purification step of the N-terminally His_6_-tagged Rlig1 was performed via a 5 mL HisTrap^TM^ FF crude column (Merck) (Buffer A: 50 mM Tris-HCl pH 8.0, 150 mM NaCl, 1.0 mM DTT, and 20 mM imidazole; Buffer B: 50 mM Tris-HCl pH 8.0, 150 mM NaCl, 1.0 mM DTT, and 500 mM imidazole). Fractions containing Rlig1 were pooled and dialysed at 4°C overnight against Buffer C (50 mM Tris-HCl pH 8.0, 100 mM NaCl, and 1.0 mM DTT). Pure Rlig1 was eluted from HiTrap® Q HP column (Merck) with Buffer D (50 mM Tris-HCl, pH 8.0, 1000 mM NaCl, and 1.0 mM DTT). After concentration, the resulting recombinant protein was transferred to storage buffer (50 mM Tris-HCl pH 8.0, 100 mM NaCl, 1 mM DTT and 50% (v/v) glycerol) and stored at −20°C.

### 5′ ^32^P-labelling of oligonucleotides

Oligonucleotides used as substrates for the ligation assay are the same as in the work of Yuan *et al.* with the human Rlig1 protein^4^. DNA and RNA oligonucleotides (1.0 μM) were incubated with 15 units of T4 PNK (New England BioLabs, M0202S) and 200 μM 0.555 MBq γ-32P-ATP (185 TBq/mmol, Hartmann Analytic, SRP-401) respectively in 1× T4 PNK reaction buffer at 37°C for 1 h. The total reaction volume amounted to 15 μL. Heating to 95°C for 2 min stopped the labelling reaction. Gel filtration using SephadexTM G-10 resin was used to remove excessive γ-32P-ATP. This led to 5′ 32P-labelled oligonucleotides of 1.0 μM.

### RNA ligation assays

To study the RNA ligation activity of zebrafish Rlig1, 0.1 μM 5′ ^32^P-labelled oligonucleotide substrate and 0.5 µM unlabelled oligonucleotide substrate were incubated with 1 μM zebrafish Rlig1 and 200 μM ATP in the RNA ligation buffer (50 mM Tris-HOAc pH 7.0, 5 mM MgCl_2_, and 1 mM DTT) at 37°C in a total volume of 10 μL. Aliquots (1.5 μL) were taken at 0, 5, 10, 30, and 60 min and quenched by adding 28.5 μL stopping solution (80% (v/v) formamide, 20 mM EDTA, 0.025% (w/v) bromophenol blue, and 0.025% (w/v) xylene cyanol) and heating to 95°C for 2 min. The resulting mixture was further diluted to 0.005 μM 5′ ^32^P-labelled oligonucleotides with stopping solution. 1 μL of the mixture was resolved by urea-PAGE (12%). Analysis of the samples was realised by autoradiographic imaging (Typhoon™ FLA 9500 biomolecular imager, GE Healthcare).

### Analysis of RNA product formation

The percentage of product formation was determined using Image Lab (Version 6.1.0). The program allows easy identification of lanes and bands. In order to calculate product formation, all visible bands were taken into consideration: (circularised) educts, products, and degradation fragments. Intensity readout computed by Image Lab directly gives the amount of product formation. The experiments were conducted in biological replicates.

### Zebrafish husbandry

Zebrafish husbandry and research was performed in accordance with the guidelines of the EU directive 2010/63/EU, the German Animal Welfare Act and the State of Baden-Württemberg (Germany) and approved by the Regierungspräsidium Freiburg. Adult zebrafish were maintained under standard conditions^53^. Embryos were incubated in embryo medium (250 mg/L Instant Ocean salt, 1 mg/L methylene blue in reverse osmosis water adjusted to pH 7 with NaHCO_3_) at 28°C unless otherwise noted. Maternal-zygotic homozygous and heterozygous *rlig1* mutants were generated by a CRISPR/Cas9-mediated knockout of the *zgc:103499* gene (animal experiment permits G-21-125 and G-24-092).

### Generation of a CRISPR rlig1^-/-^ knockout fish line

To generate a zebrafish *rlig1* null mutant that does not produce any *rlig1* mRNA, the promoter region of the *rlig1* gene in zebrafish (*zgc:103499*) was deleted using the CRISPR/Cas9 system. The decision to remove all possible transcriptional start sites is based on the observation that truncated mRNA transcripts may induce transcriptional adaptation. This could lead to the up-regulation of related genes with similar functions^54^, which in turn could serve to compensate for the *rlig1* knockout^55^. To this end, the transcription start sites of *rlig1* were mapped using CAGE data along with H3Kme3 and RNA-Seq data from Nepal *et al.*^56^. The upstream genomic sequence showing CAGE signals in any of the embryonic stages – covering the entire first exon of *rlig1* and part of the second exon, which was annotated to contain an alternative transcription start site (NC_007136.6, Annotation Software Version 7.4) – were targeted for deletion (in total ∼810 bp).

Two crRNAs, one targeting a sequence directly upstream of the identified promoter region and one targeting the second intron of *rlig1* (Table 1), were obtained from IDT. Guide RNAs (gRNAs) were prepared as described by Kroll *et al.*^57^ (see https://doi.org/10.17504/protocols.io.bs2rngd6). In short, crRNAs were annealed with a trans-activating crRNA (Alt-R CRISPR-Cas9 tracrRNA, IDT, 1072532), and the resulting gRNAs were assembled separately with the Cas9 protein (Alt-R S.p. Cas9 Nuclease V3, IDT, 1081058). After pooling the two RNPs in equal amounts, 1 nL of the RNP mix (19 µM) was injected into the yolk of early 1-cell stage zebrafish embryos. The efficiency of the CRISPR-mediated knockout was determined by PCR^58,59^ (Supplementary Figure 2).

**Table 1.**
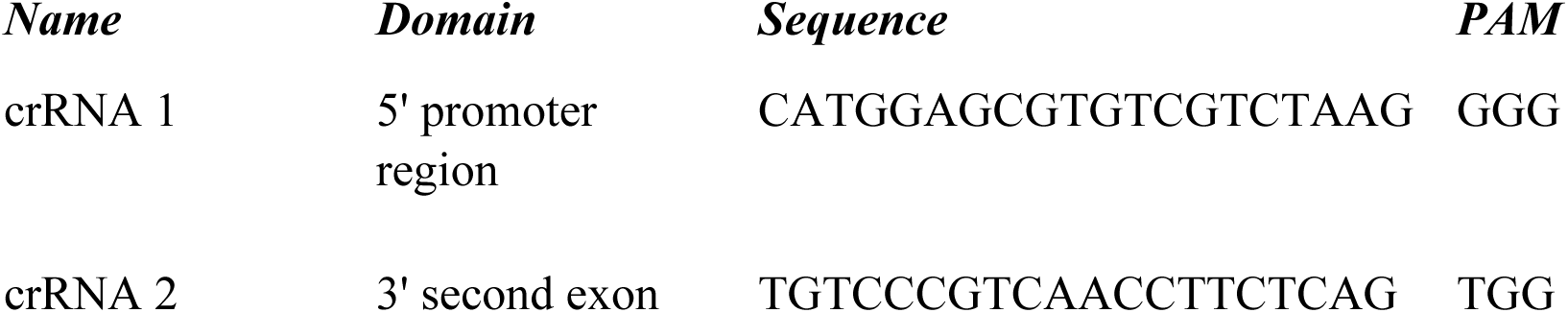
CRISPR RNAs (crRNAs) targeting *rlig1*.

**Table 2.**
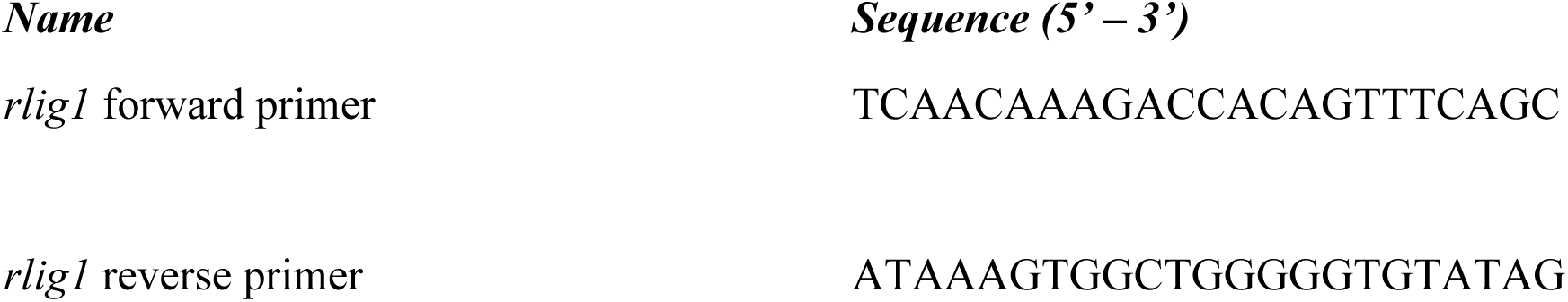
Sequences of primers used for genotyping the *rlig1*^-/-^ allele.

### Genotyping

PCR was performed to evaluate the cutting efficiency of RNPs and verify the successful knockout of the *zgc:103499* promoter region in the P and all following F generations. For this purpose, genomic DNA was isolated either from a pool of 10 P *rlig*^-/-^ embryos 1 dpf or by fin-clipping of adult individuals using the “hotshot” method^60^. The PCR was conducted under standard PCR conditions, leading to the amplification of a sequence spanning the targeted promoter region. Upon successful deletion (KO), a 334 bp long product is amplified, whereas unsuccessful cleavage or remaining of at least one WT allele yields a 1148 bp long product. We used the *rlig1^kn101pm^* mutant allele (Supplementary Figure 2) for all experiments and analyses presented here.

### Total RNA extraction

To investigate *rlig1* mRNA levels during embryonic development, 10-20 embryos of the desired developmental stage were pooled, snap-frozen and stored at −80°C until further use. Total RNA was obtained by homogenising the embryos with a pestle and subsequent extraction of RNA using the TRIzol™ Reagent (Invitrogen) following the manufacturer’s guidelines. In short, embryos were lysed in TRIzol™ Reagent (Invitrogen), followed by RNA precipitation with isopropanol (overnight, −80°C). Washing of the obtained RNA pellet with ethanol removed remaining salts. After resuspending the RNA in RNase-free water, potential DNA contaminations in the sample were digested by incubation with DNase I-XT (NEB) for 30 min at 37°C. Further purification was conducted following the manual instructions of the RNA Clean & Concentrator-25 (ZYMO RESEARCH) Kit. RNA quality and concentration was determined with the Bioanalyzer System (Agilent). The RNA was stored at −80°C.

### Reverse transcription quantitative polymerase chain reaction (RT-qPCR)

The elongation factor 1 alpha (*eef1a)* was chosen as a reference gene due to its stable expression during embryonic development for later normalisation across samples^15^. TaqMan probes for *rlig1* and *eef1α* were designed to target regions spanning over two exons and ordered from IDT (Table 3). The *rlig1* probe was designed bearing a 5’ Cy5^TM^ dye and a 3’ Iowa Black® RQ quencher. The housekeeping probe employed the 5’ SUN™ dye and 3’ Iowa Black® FQ as a quencher.

**Table 3.**
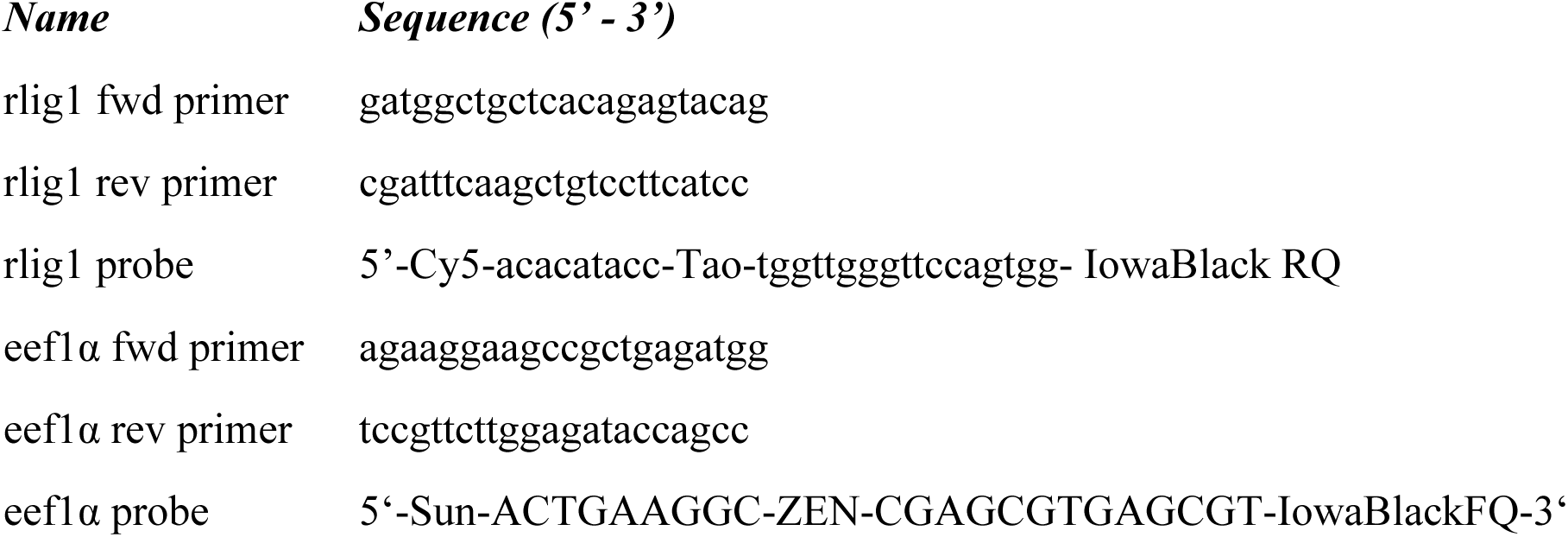
Primer and probe sequences.

In each RT-qPCR reaction, 100 ng extracted total RNA served as a template. The Luna® Universal Probe One-Step RT-qPCR Kit (NEB) was used for optimal amplification of the target sequences. 1x Luna® Universal Probe One-Step Reaction Mix and 1x Luna® WarmStart® RT Enzyme Mix was used for qPCR with 0.67 µM of the respective forward and reverse primers (Table 3) and 0.17 µM of the respective probes in 10 µL reaction volume. The initial reverse transcriptase step at 55°C for 10 min was followed by an initial denaturation step at 95°C for 1 min. Next, a two-step cycling protocol with 45 PCR cycles of a denaturation step at 95°C for 10 s and an extension step at 64.1°C for 60 s was employed. Fluorescent intensity of both probes was measured in the LightCycler® 96 System (Roche) at the end of each cycle. The LightCycler^®^ 96 Application and Instrument Software was used for data visualisation and analysis. To ensure reproducibility, the experiment was conducted in biological triplicates. For the purpose of comparability of different samples, the obtained Cq value for the *rlig1* probe was normalised by subtracting the Cq value of *eef1α* in the respective sample, yielding a normalised ΔCq value for each sample. GraphPad Prism 6 was employed to combine the measurements in triplicates and present the obtained data in one graph. Plotting the reverse ΔCq value (1/ΔCq) on the *y*-axis enabled the depiction of the relative levels of *rlig1* mRNA during selected embryonic developmental stages.

### HCR RNA FISH

Embryos of the desired developmental stages (2-8 cell stage, sphere, shield, 1 dpf, 2 dpf, 3 dpf, and 5 dpf) were fixed in 4% paraformaldehyde over night at 4°C. For storage and permeabilisation, the embryos were stepwise transferred to methanol and stored at −20°C. Stepwise rehydration of the embryos to PBST (1× PBS + 0.1% Tween-20) was followed by dechorionation (0-1 dpf) or depigmentation (2-5 dpf). Depigmentation was achieved by incubation of the embryos with 3 % H_2_O_2_ solution in KOH under an LED light source. All HCR reagents were purchased from Molecular Instruments®. The number of embryos per 2 mL tube depended on the age: up to 14 embryos for embryos < 1 dpf, 12 embryos for 1 dpf, 10 embryos for 2 dpf, and 8 embryos for 3 dpf and 5 dpf. The protocol from the Baier lab^19^ was followed for the HCR RNA FISH procedure. First, the embryos were pre-hybridised with probe hybridisation buffer (30 min, 37°C). This was followed by hybridisation with the *rlig1* probe set (LOT RTJ881, 2 pmol probe set, 16 h, 37°C). Excessive, unbound probe was removed by washing the embryos four times with probe wash buffer (15 min, 37°C). Subsequently, the embryos were transferred to 5x SSC-T at room temperature (RT) until the amplification step was initiated. The addition of amplification buffer (30 min, RT) set a pre-amplification environment. In the meantime, the fluorescently labelled hairpins were prepared by heating to 95°C for 90 s and subsequent cooling to RT for 30 min in the dark. The hairpin solution was prepared by adding 30 pmol of each hairpin (B2-546/ B2-647) to the amplification buffer. The amplification step occurred overnight in the dark (16 h, RT). Excessive unbound hairpins were removed by washing three times with 5x SSC-T (20 min, RT) in the dark. Samples that required further staining of the nucleus were incubated with DRAQ7™ Dye (Thermo Fisher) diluted 1:1000 in 5x SSC-T. Incubation times differed between the developmental stages with 30 min at RT for 2-8 cell stage embryos, sphere embryos and shield embryos, 5 h at RT for 1 dpf old embryos, and 8 h at RT for 2-5 dpf old embryos. Excessive DRAQ was removed by washing three times with 5x SSC-T (15 min, RT). Larvae that were selected for high-resolution imaging with the Zeiss LSM 880 microscope were stained with DAPI (Sigma, D9542) diluted 1:500 in 5x SSC-T. The incubation period amounted to 3 days at 4°C in the dark. Excessive DAPI was removed by washing three times with 5x SSC-T (15 min, RT). Storage until imaging occurred in 5x SSC-T at 4°C in the dark.

### Imaging of fluorescence in situ hybridisation samples

Fluorescence images of *rlig1 in situ*-stained embryos were obtained using a Zeiss Lightsheet Z.1 selective plane illumination microscope (SPIM) equipped with a W Plan-Apochromat 10× objective. Embryos were mounted in 1.5% low-melting agarose (Lonza 50080) in a size 3 glass capillary (Zeiss) and orientated with the help of a dissecting needle. Filters and light sheets were auto-aligned prior to imaging. *rlig1* mRNA (546 nm-amplifier fluorophore, see above) and DRAQ7 were imaged sequentially: *rlig1* mRNA using a 561 nm laser (20% intensity) with a BP 575-615 filter and 100 ms exposure and DRAQ7-stained nuclei using a 638 nm laser (10% intensity) with LP 660 filter and 70 ms exposure. z-stacks of 306 – 630 µm (with 1.8164 µm interval between z slices) were taken from two orientations, lateral views and animal-pole or dorsal views. Zoom settings altered depending on the embryo stage with 0.85-1.0 x for embryos < 1 dpf (cleavage: 0.9x lateral and 1.0x animal pole, sphere: 0.85x, shield: 1.0x) and 0.8x for embryos ≥ 1dpf. Likewise, the number of tiles imaged was dependent on embryo stage (<1 dpf: 1 tile, 1 dpf: 1-2 tiles, 3 dpf: 4 tiles, 5 dpf: 6 tiles) as well as of the mounting orientation within the glass capillary. Stitched images as well as maximum intensity projections were generated using ImageJ 1.54f software^61^ and used for the analysis and visualisation of the data.

To gain high-resolution images of the *rlig1* mRNA signal in the brain at 5 dpf, MZ*rlig1* and WT larvae were embedded upside down in 1.5% low melting agarose in a CELLview cell culture dish (4 compartments, Greiner-Bio, #627975). Imaging of the larval heads was performed on a Zeiss LSM 880 Airyscan microscope equipped with a 25x multi-immersion objective (LD LCI Plan-Apochromat 25x/0,8 Imm Korr DIC M27, Zeiss, catalog no. 420852-9871-000) using water immersion and a 0.7x zoom. *rlig1* mRNA (647 nm-amplifier fluorophore, see above) and DAPI were imaged sequentially using a 633 nm and a 405 nm laser with an intensity of 55% and 1.1% and a gain of 780 and 770, respectively. *z*-stacks, comprising two tiles covering the entire brain, were taken with a pixel dwell time of 0.53 µs. Pre-processing of imaging data in ZEN 3.0 black (Zeiss) and subsequent stitching in ZEN 3.0 blue (Zeiss) generated a final image with a size of 3694 x 1958 pixels (922.03 x 493.11 µm, 0.492 µm in z) for MZ*rlig1* larvae and 3694 x 1958 pixels (911.91 x 483.36 µm, 0.492 µm in z) for WT larvae. Maximum intensity projections were generated using ImageJ 1.54f software^61^ and used for the analysis and visualisation of the data.

### Image analysis

The ImageJ 1.54f software was used for image stitching and analysis. To visualise the distribution as well as the intensity of fluorescent signal, either singles slices were selected or z-projections spanning over multiple slices were generated displaying the maximum intensity. The brightness and contrast settings for the *rlig1* mRNA signal were set identically in the captured images of WT and MZ*rlig1* embryos of the same developmental stage. Conversely, the brightness and contrast parameters for the DRAQ7 or DAPI reference channels were modified to produce images of comparable intensity, regardless of their actual intensity.

### Behaviour experiments

Behavioural experiments were conducted with 5 dpf old freely swimming larvae of WT or MZ*rlig1* genotype. To minimise genomic variability between the two groups, cousin controls were used, meaning that the parents of the larvae were siblings. The embryos were raised in small groups (30-40 individuals) in Petri dishes (14 cm in diameter) on a 14 h light, 10 h dark cycle at a constant 28°C temperature. Closed-loop behaviour paradigms were performed on a system adapted from our previous work^28^. To investigate the behaviour of 5 dpf old larvae, individuals were transferred into custom-designed circular acrylic dishes (12 cm in diameter, 5 mm in height, black rim, transparent base covered with diffusion paper) filled with 50 mL of E3 fish water. Visual stimuli were projected onto the dish from below with AAXA Neo-pico projectors and the dishes were illuminated for the camera using infrared light-emitting diode (LED) strip panels (940 nm panel, SOLAROX®). A high-speed CMOS camera (Basler acA2040-90um-NIR or Grasshopper3-NIR, FLIR Systems) with a zoom lens (#58-240-6X, 18-108mm FL, Edmund Optics) and an infrared filter (850 nm) were used to track larvae. Live tracking of the larvae was realised in real-time at a rate of 90 fps using custom-written behavioural tracking software on Python 3.12. The larval position was determined from the largest centre-of-mass after background subtraction. Additionally, second-order image moments were used to define larval orientation. In order to identify events of high activity (bouts), a 50 ms rolling variance window over body orientation was calculated. Swim bouts were defined to start once the rolling window variance exceeded 1 deg^2^ for at least 20 ms. Swim bouts ended once the variance dropped below 0.5 deg^2^ for at least 50 ms. High throughput was achieved by tracking up to 8 larvae simultaneously in a single setup. This was made possible by connecting one computer to 8 cameras and 4 projectors covering 8 dishes.

### Visual stimulus setup

The visual stimuli for the behaviour experiments consisted of moving gratings (speed: 1.8 cm/s, wavelength 0.6 cm) and grey uniform control. Unless mentioned otherwise, each trial contained 10 s grey, 20 s stimulus, 15 s grey. Grating directions (left, right, forward, grey control) were presented in random order.

The visual stimuli for the imaging experiments consisted of random dot motion kinematograms. The motion stimulus parameters (1.8 cm/s, 100% coherence) were selected based on previous studies^28^ showing robust activation of direction-selective neurons in the pretectum and anterior hindbrain of larval zebrafish. In short, ∼1000 dots (2 mm in diameter) with a lifetime of (200 ms mean) stochastically disappeared and immediately reappeared at a random location. During left or right stimuli, 100% of the dots moved at 1.8 cm/s. Each trial contained 10 s 0% coherence (no overall movement), 30 s 100% coherence, 10 s 0% coherence. Dot directions (left, right, 0% coherence control) were presented in random order.

### Behaviour analysis

Behaviour data was analysed at the swim bout level. Orientation change, interbout interval (time between two bouts), and distance per bout were extracted both during left/right stimulus and during the grey control. Next, the bouts were binarised into left and right (ignoring bouts between −2° and 2°). A 2 s rolling average/SEM window was calculated to obtain the percentage left over trial time graph.

### Two-photon calcium imaging

For the purpose of investigating neuronal activity of WT and MZ*rlig1* zebrafish larvae, MZ*rlig1* fish were crossed with Tg(*elavl3*:*H2B-GCaMP8s)^mpn43830^* fish carrying a *mitfa(nacre)* background. The *mitfa* mutants used in this study originate from a background containing the *mitfa^s170^* and *mitfa^s184^* alleles^62^; the mutants lack melanocytes, thereby facilitating imaging of the genetically encoded calcium sensor GCaMP8s. 5 dpf old larvae were screened and selected for strong GCaMP8s fluorescence and no pigmentation. The larvae were embedded in 2% agarose (UltraPure Low Melting Point Agarose, 16520-100, Invitrogen) in a small Petri dish (6 cm in diameter) and imaged in a custom-built two-photon microscope, operated by custom-written Python 3.12-based software (PyZebraPhysiology). Briefly, a femtosecond-pulsed MaiTai Ti-Sapphire laser (Spectra Physics) tuned to 950 nm, a set of x/y-galvanometers (Cambridge Technology), and a 25x objective (MRD77220, Nikon) were used to scan over the brain. A sensitive GaAsP alkali photomultiplier tube (Hamamatsu), amplified by a current preamplifier (TIA60, Thorlabs) was used to record GCaMP fluorophore emission. Frame acquisition rates of around 1 Hz were used in all imaging experiments, and the laser power was adjusted to ∼12 mW at the specimen. To image neuronal activity in the anterior hindbrain, tectum (midbrain) or pre-tectum (forebrain), a single plane (spatial resolution: 0.28 µm per pixel, 800×800 pixels) was imaged for 1 h. The stimuli were projected through a mirror by a P300 Neo Pico projector (Aaxa Technologies) onto diffusive paper glued to the bottom of the experimental platform (8 cm diameter). To prevent accidental bleed-through of non-red light into the photomultiplier, the projector was prepared with a red foil in front of the lens. An overview stack of most of the brain was imaged after each imaging session by recording ∼25 planes at 5 µm distance (spatial resolution: 0.56 µm per pixel, 800 x 800 pixels) for 20 s per plane. The overview stack was used as a reference for anatomical mapping of the functional plane.

### Pre-processing two-photon calcium imaging data

The raw imaging data was pre-processed by a custom-written Python 3.10-based script. In brief, motion-correction, manual region segmentation, automatic cell segmentation, stimulus temporal alignment and anatomical registration were performed. Piece-wise rigid motion correction (*NoRMCorre*^63^) was performed using the *CaImAn*^64^ framework and standard parameters. Next, in each hemisphere of the imaged brain structures, the region containing direction-selective cells was manually outlined. This was followed by using Cellpose^31^ to automatically segment single cells using [model_type = ‘cyto3’^65^, cellpose_flow_threshold = 0.4, cellpose_prob_threshold = 0] and otherwise default parameters. Thereafter, the average fluorescence of each region and cell was temporally aligned using SciPy’s *interp1d* function^66^ with a discretisation of 0.5 s. Based on the segmented masks, single-cell or region-wide calcium dynamics from the stimulus-aligned data were extracted. Calculation of dF/F0 = (*F* − *F*₀) / *F*₀ for each segmented cell/region was done by taking the 5^th^ percentile of F as the F₀ baseline. Finally, a two-step registration strategy based on the image registration library ANTs^67^ was used. In the first step, the overview stack was mapped to the ZBRAIN^68^, and in the second step, the functional plane was mapped through the overview stack to the ZBRAIN coordinates.

### Analysis and statistics of two-photon calcium imaging data

Region-wide responses in the anterior hindbrain were investigated by examining the dF/F0 trial traces of each region during the last 20 s of the preferred motion stimulus (i.e., leftward motion for the left-hemisphere, rightward motion for the right hemisphere). The statistical significance of the difference in response of WT and MZ*rlig1* fish was calculated using a one-sided t-test.

Next, each segmented cell was tested for a response to motion. A t-test was performed between the peak dF/F0 of the pre-stimulus baseline and that of the response periods. Only neurons that showed a significant increase during stimulus (p < 0.005) were considered motion-responsive. The direction selectivity index (DSI) was calculated as follows:

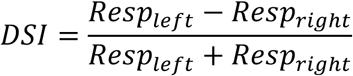

A Gaussian was fitted to the overall DSI distribution of WT and MZ*rlig*1 combined, and 2*STD of this distribution was used as the threshold to define direction-selective neurons (abs(DSI) > 0.273). Finally, the ratio of direction-selective neurons among the motion-responsive neurons was calculated. The analysis was performed separately for neurons located in the forebrain (mostly pretectum), midbrain (mostly tectum), and hindbrain (mostly anterior hindbrain). If less than 100 neurons per plane were found for a region, this plane/region combination was ignored. Statistical testing was performed separately for each brain region. To reduce the risk of inflated false-positive rates due to multiple comparisons, statistical comparisons across brain regions were subsequently conducted using one-sided t-tests with Bonferroni correction.

### Sample preparation for mRNA sequencing

Bulk sequencing analysis was performed to identify transcriptomic differences between WT and MZ*rlig1* larvae. For developmental profiling, samples were collected at six stages: 4-cell, sphere, shield, bud, 1 dpf, and 5 dpf. Per condition, 10 embryos were pooled for the 4-cell, sphere, and shield stages, and 8 embryos for the bud, 1 dpf, and 5 dpf stages. Pooled embryos were snap-frozen in liquid nitrogen and stored at –80 °C. Total RNA was extracted using TRIzol™ Reagent (Invitrogen) according to the manufacturer’s instructions. RNA integrity was assessed using the Bioanalyzer System (Agilent Technologies). In addition, a separate transcriptome analysis focused specifically on the heads of the larvae, taking their individual behavioural performance into account. To be categorised as “good” performers, larvae had to meet two criteria: (A) executing overall more than 50 bouts following the stimulus and (B) performing more bouts following the stimulus than in the opposite direction. Leftward swims were defined as orientation changes of ≥ 28.8°, while rightward swims were defined as orientation changes of ≤ −28.8° relative to the larval body axis. Larvae that either failed to reach the 50-bout threshold or did not predominantly swim in the stimulus direction were classified as “bad” performers. To ensure accurate behavioural classification, larvae were exposed to alternating leftward and rightward grating stimuli for a total duration of 1 h. Immediately after the assay, larvae were anesthetised on ice, and heads were dissected using a scalpel. For each sample, heads from eight larvae of identical genotype and behavioural classification were pooled, snap-frozen in liquid nitrogen, and stored at –80 °C. Total RNA was prepared and quality-checked as described above.

### mRNA sequencing and differential gene expression analysis

Library preparation, quality control, and sequencing were carried out by Novogene Bioinformatics Technology Co., Ltd. (Munich, Germany). mRNA sequencing was performed on the Illumina NovaSeq X Plus platform, generating 150 bp paired-end reads with a sequencing depth of approximately 20 million reads (∼6 Gb) per sample. Library preparation followed Novogene’s standard protocol, which included poly-T oligo-attached magnetic bead–based mRNA enrichment, fragmentation, first- and second-strand cDNA synthesis, end repair, A-tailing, adapter ligation, size selection, PCR amplification, and final purification.

For the developmental time-course experiment, samples were sequenced for both WT and MZ*rlig1* embryos at the 4-cell stage (quadruplicates, n = 40 per genotype), sphere stage (duplicates, n = 20 per genotype), shield stage (triplicates, n = 30 per genotype), bud stage (triplicates, n = 24 per genotype), 1 dpf (triplicates, n = 24 per genotype), and 5 dpf (triplicates, n = 24 per genotype). A total of 13 samples were sequenced for the head-based behavioural dataset, comprising four WT-good (n = 32), three WT-bad (n = 24), two MZ*rlig1*-good (n = 16), and four MZ*rlig1*-bad (n = 32) biological replicates.

Bioinformatic processing started with the download of the reference genome (Danio_rerio.GRCz11.dna_sm.primary_assembly.fa.gz) and gene annotation file (Danio_rerio.GRCz11.113.gtf.gz) from ENSEMBL (release 113)^69^. An index was generated using *STAR* (v.2.7.10b)^70^, and clean paired-end reads were then aligned to the reference genome using the same version of *STAR*. To reconstruct the transcriptome, the genome FASTA and annotation GTF files were processed with *gffread* (v.0.12.7)^71^. Transcript- and gene-level expression quantification was subsequently performed with *Salmon* (v.0.14.1)^72^ in alignment-based mode, using *STAR*-generated transcriptome BAM files as input.

Differential gene expression (DGE) analysis was conducted in R using the *DESeq2* package (v.1.42.1)^73^. Transcript-level quantifications from *Salmon* were imported with the *tximport* package (v.1.30.0)^74^, and a DESeqDataSet object was created from the count data and corresponding experimental metadata, with ‘condition’ defined as the main factor of interest. Genes with fewer than 10 reads across all samples were excluded to reduce noise. Counts were normalised using DESeq2’s median-of-ratios method, and p-values were adjusted for multiple testing using the Benjamini–Hochberg false discovery rate (FDR). Genes with an adjusted p-value (padj) < 0.05 were considered differentially expressed.

For read coverage analysis, BAM files were sorted and indexed using *samtools* (v.1.13)^75^. Gene ontology (GO) and KEGG pathway enrichment analyses were performed with *clusterProfiler* (v.4.10.1)^76^. GO enrichment was carried out separately for upregulated and downregulated gene sets, assigning significant terms to the categories of molecular function (MF), cellular component (CC), and biological process (BP). KEGG analysis was likewise conducted for both up- and downregulated genes to identify enriched biological pathways. In addition, gene set enrichment analysis (GSEA) was performed using *clusterProfiler* to identify consistently enriched pathways across the entire ranked gene list.

