## Supplementary Information for "An RNA ligase shapes transcriptional profiles, neural function, and behaviour in the developing larval zebrafish"

**Contents**

Supplementary Figures 1-5

Supplementary Tables 1-14

Supplementary Movies 1-3

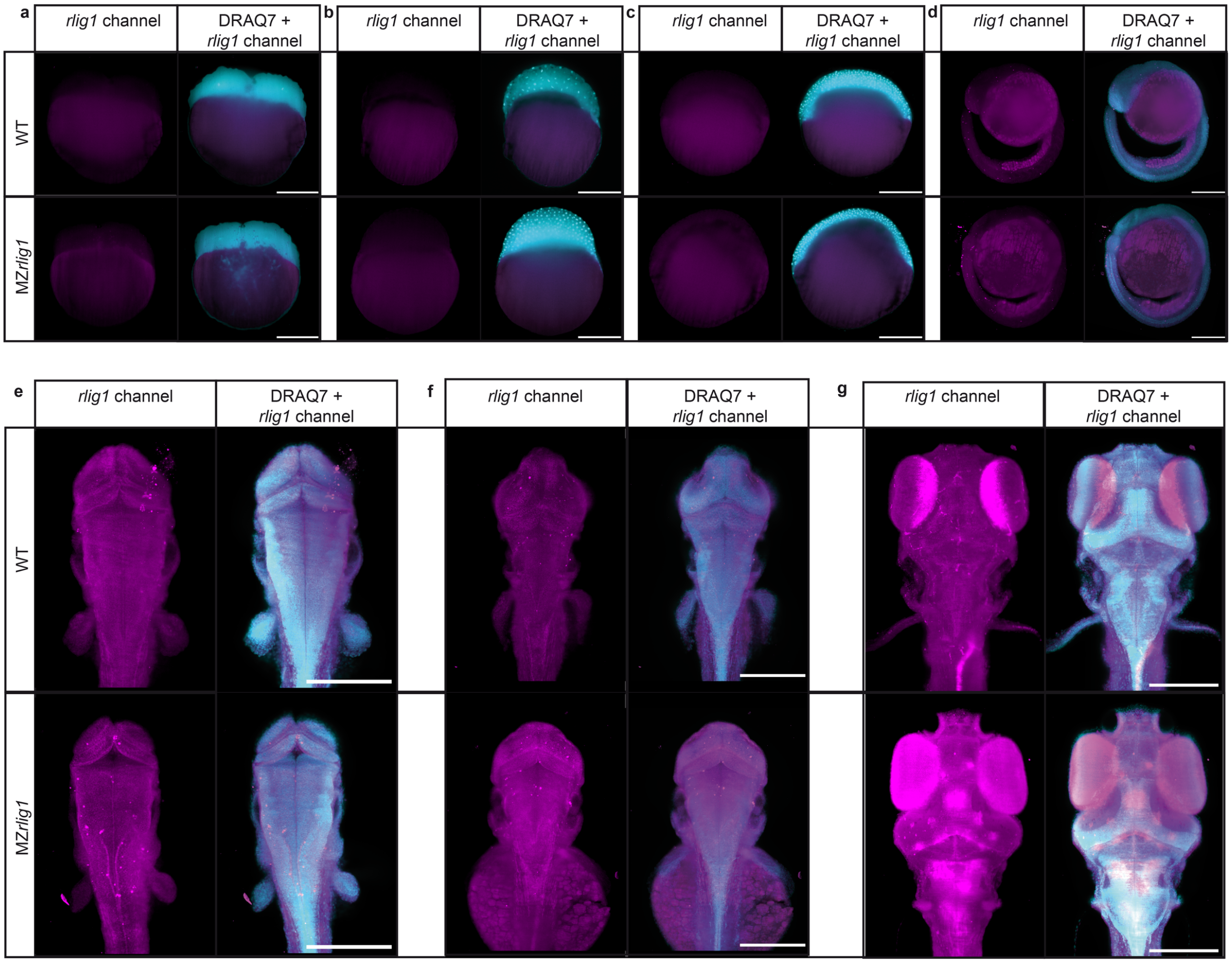

**Supplementary Figure 1 | Background signal assessment for HCR RNA FISH. (a–g)** Assessment of background signal by imaging in the *rlig1* channel without *rlig1* probe application. The *rlig1* detection channel is shown in magenta, the DRAQ7 reference in cyan. Seven developmental stages were selected for imaging of WT and maternal-zygotic *rlig1*$^{-/-}$ mutants (MZ*rlig1*). **(a)** 4-cell stage, lateral view. **(b)** Sphere stage, lateral view. **(c)** Shield stage, lateral view. **(d)** 1 dpf, lateral view with anterior to the left. **(e)** 2 dpf, dorsal view of the trunk region with anterior to the top. **(f)** 3 dpf, dorsal view of the trunk region with anterior to the top. **(g)** 5 dpf, dorsal view of the trunk region with anterior to the top. (a–c) show single optical slices, while (d-g) show maximum intensity projections. Scale bars: 250 μm.

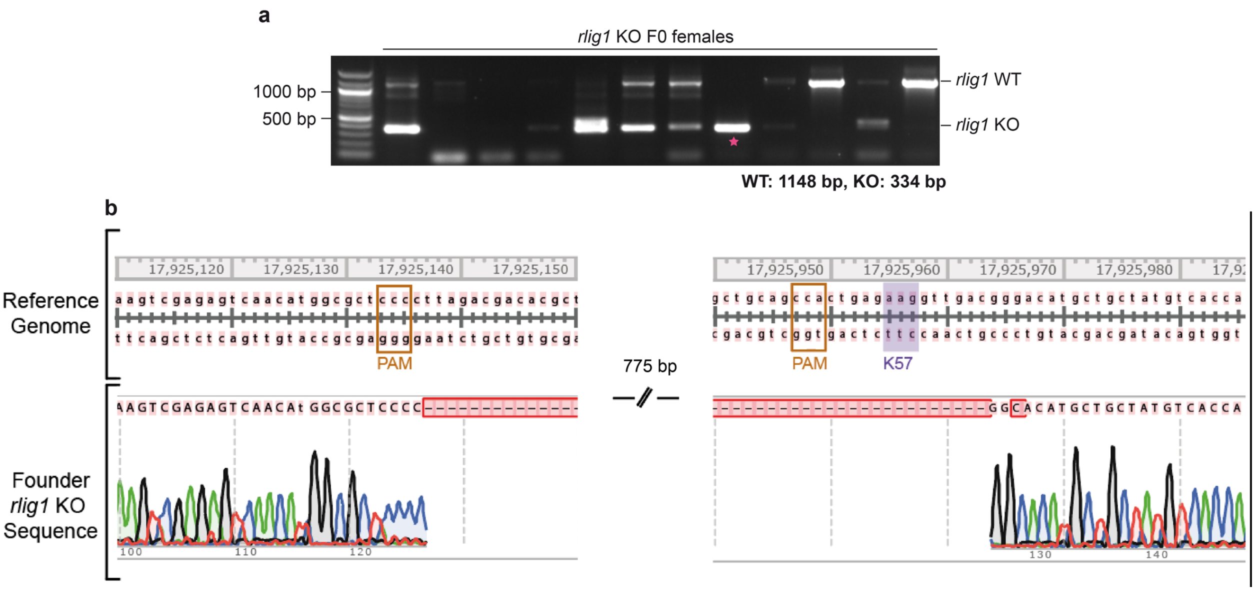

**Supplementary Figure 2 | Generation of zebrafish *rlig1* knockout animals. (a)** Validation of the *rlig1* knockout on the genomic level by PCR. Amplification of genomic sequence spanning the *rlig1* deletion site. The WT allele produces a fragment of 1148 bp, whereas the deleted allele yields a 334 bp product. The pink asterisk indicates the founder female that was used for establishing the MZ*rlig1*$^{kn101pm}$ line. **(b)** Genomic sequence of the *rlig1* founder female determined by Sanger sequencing. The upper row shows the reference sequence and genomic positions (GRCz11, accession NC_007136), with the PAM sites of the two gRNAs used for deletion highlighted in light brown and the codon for the catalytically relevant lysine residue (K57) indicated by a purple shaded area. The lower row shows the genomic sequence of the founder female, with the deletion sequence highlighted in red.

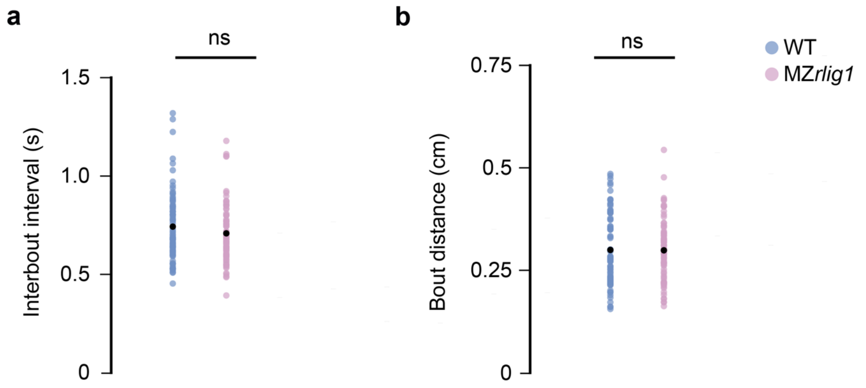

**Supplementary Figure 3 | Analysis of interbout interval and bout distance in WT and MZ*rlig1* larvae under no-motion stimulus conditions. (a)** Interbout interval (s) of individual larvae in the no-motion condition (grey uniform control stimulus). Each dot represents the mean value of a single larva, averaged across all trials. WT larvae are shown in blue, MZ*rlig1* larvae in pink. Black dots indicate the group mean. Mann–Whitney U test: ns = non-significant. **(b)** Bout distance (cm) under the same condition. Data presentation and colouring as in (a). Mann–Whitney U test: ns = non-significant. Representative data from n = 80 WT and MZ*rlig1* larvae.

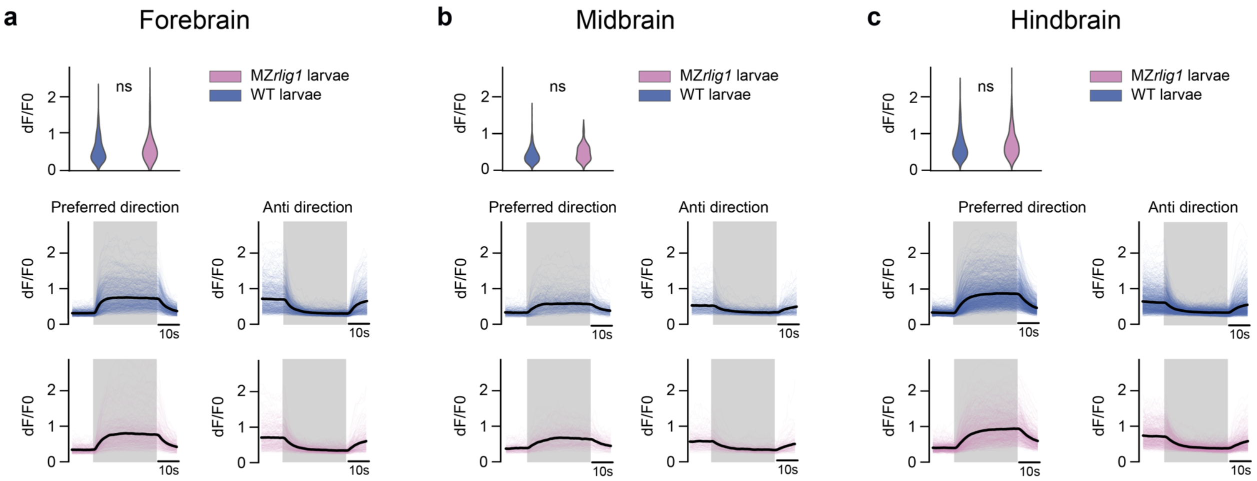

**Supplementary Figure 4 | Neuronal activity of direction-selective neurons in forebrain, midbrain, and hindbrain of WT and MZ*rlig1* larvae.** Upper plots: Mean dF/F0 activity of direction-selective neurons in WT (blue) and MZ*rlig1* (pink) larvae in **(a)** forebrain, **(b)** midbrain, and **(c)** hindbrain. Statistical comparison based on the mean dF/F0 during 20–40 s (one-sided t-test); ns = non-significant. Lower plots: Temporal dynamics of neuronal activity in the same brain regions. Blue: WT, pink: MZ*rlig1*. Individual traces across trials are shown with the mean in black. Preferred and anti-direction are indicated; stimulus duration is shaded in grey. Neuronal activity is represented as dF/F0. Sample size: MZ*rlig1* hindbrain: 12 planes from 11 larvae, midbrain: 18 planes from 17 larvae, forebrain: 10 planes from 10 larvae; WT hindbrain: 14 planes from 12 larvae, midbrain: 22 planes from 20 larvae, forebrain: 13 planes from 11 larvae.

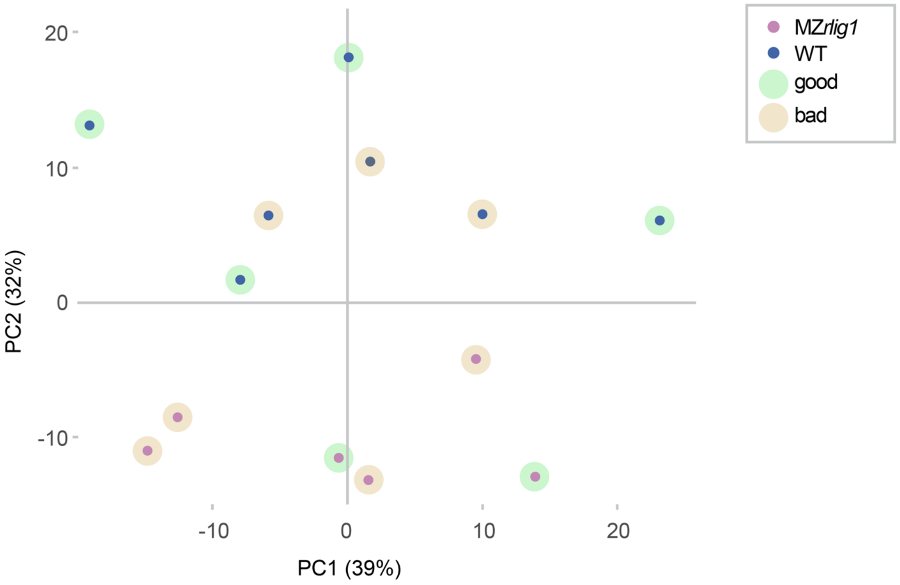

**Supplementary Figure 5 | Principal component analysis (PCA) of larval head transcriptomes based on gene expression values (FPKM).** WT (blue) and MZ*rlig1* (pink) samples are shown, with behavioural classification indicated by shaded backgrounds: green, “good” responders; orange, “bad” responders.

**Supplementary Table 1 | Differential expression of *rlig1* across developmental stages of WT embryos.** Normalised expression data for *rlig1* obtained by RNA-Seq at six developmental stages. The table includes average expression levels (baseMean), $log_2$FoldChange (MZ*rlig1* vs. WT), and adjusted p-values (padj) from DESeq2 analysis.

| Stage | baseMean | $log_2$(FC) | padj |
|---|---|---|---|
| **4-cell** | 436.18 | -10.89 | 6.05 E-23 |
| **sphere** | 132.72 | -10.81 | 1.28 E-11 |
| **shield** | 13.11 | -7.03 | 5.05 E-05 |
| **bud** | 23.88 | -8.30 | 2.21 E-08 |
| **1 dpf** | 60.54 | -8.96 | 3.09 E-07 |
| **5 dpf** | 95.07 | -9.59 | 1.87 E-12 |

**Supplementary Table 2 | Functional enrichment analysis for the 4-cell stage (MZ*rlig1* compared to WT).** Listed are significantly (padj < 0.05) enriched terms from Gene Ontology (GO) subdivided into biological process (BP), cellular component (CC), and molecular function (MF), gene set enrichment analysis (GSEA) terms, and KEGG pathways.

| Term | ID | Description | padj | q-value |
|---|---|---|---|---|
| **GO-BP** | GO:0006119 | oxidative phosphorylation | 0.000589 | 0.000556 |
| **GO-BP** | GO:0042773 | ATP synthesis coupled electron transport | 0.000589 | 0.000556 |
| **GO-BP** | GO:0009060 | aerobic respiration | 0.000589 | 0.000556 |
| **GO-BP** | GO:0022904 | respiratory electron transport chain | 0.001513 | 0.001428 |
| **GO-BP** | GO:0045333 | cellular respiration | 0.001513 | 0.001428 |
| **GO-BP** | GO:0022900 | electron transport chain | 0.001513 | 0.001428 |
| **GO-BP** | GO:0015980 | energy derivation by oxidation of organic compounds | 0.012261 | 0.011573 |
| **GO-BP** | GO:0019646 | aerobic electron transport chain | 0.013343 | 0.012594 |
| **GO-BP** | GO:0042775 | mitochondrial ATP synthesis coupled electron transport | 0.014907 | 0.01407 |
| **GO-BP** | GO:0006120 | mitochondrial electron transport, NADH to ubiquinone | 0.037129 | 0.035046 |
| **GO-CC** | GO:0098803 | respiratory chain complex | 0.001402 | 0.001187 |
| **GO-CC** | GO:1902495 | transmembrane transporter complex | 0.001402 | 0.001187 |
| **GO-CC** | GO:0070469 | respirasome | 0.001402 | 0.001187 |
| **GO-CC** | GO:1990351 | transporter complex | 0.001402 | 0.001187 |
| **GO-CC** | GO:0043235 | receptor complex | 0.017431 | 0.014766 |
| **GO-CC** | GO:0070069 | cytochrome complex | 0.018581 | 0.01574 |
| **GO-CC** | GO:0005746 | mitochondrial respirasome | 0.018581 | 0.01574 |
| **GO-CC** | GO:1990204 | oxidoreductase complex | 0.029585 | 0.025062 |

| | | | | |
|---|---|---|---|---|
| **GO-CC** | GO:0045277 | respiratory chain complex IV | 0.048653 | 0.041215 |
| **GO-CC** | GO:0098590 | plasma membrane region | 0.048653 | 0.041215 |
| **GO-CC** | GO:0005747 | mitochondrial respiratory chain complex I | 0.048653 | 0.041215 |
| **GO-CC** | GO:0030964 | NADH dehydrogenase complex | 0.048653 | 0.041215 |
| **GO-CC** | GO:0045271 | respiratory chain complex I | 0.048653 | 0.041215 |
| **GO-CC** | GO:0014069 | postsynaptic density | 0.048653 | 0.041215 |
| **GO-CC** | GO:0032279 | asymmetric synapse | 0.048653 | 0.041215 |
| **GO-CC** | GO:0098984 | neuron to neuron synapse | 0.048653 | 0.041215 |
| **GO-MF** | GO:0015453 | oxidoreduction-driven active transmembrane transporter activity | 2.76E-05 | 2.56E-05 |
| **GO-MF** | GO:0009055 | electron transfer activity | 0.00012 | 0.000112 |
| **GO-MF** | GO:0003954 | NADH dehydrogenase activity | 0.0005 | 0.000465 |
| **GO-MF** | GO:0016651 | oxidoreductase activity, acting on NAD(P)H | 0.001008 | 0.000938 |
| **GO-MF** | GO:0038023 | signalling receptor activity | 0.001075 | 0.001 |
| **GO-MF** | GO:0060089 | molecular transducer activity | 0.001075 | 0.001 |
| **GO-MF** | GO:0008137 | NADH dehydrogenase (ubiquinone) activity | 0.001382 | 0.001286 |
| **GO-MF** | GO:0050136 | NADH dehydrogenase (quinone) activity | 0.001382 | 0.001286 |
| **GO-MF** | GO:0003955 | NAD(P)H dehydrogenase (quinone) activity | 0.001752 | 0.001631 |
| **GO-MF** | GO:0015399 | primary active transmembrane transporter activity | 0.001981 | 0.001844 |
| **GO-MF** | GO:0016655 | oxidoreductase activity, acting on NAD(P)H, quinone or similar compound as acceptor | 0.001981 | 0.001844 |

| | | | | |
|---|---|---|---|---|
| **GO-MF** | GO:0004888 | transmembrane signalling receptor activity | 0.006677 | 0.006214 |
| **GO-MF** | GO:0022804 | active transmembrane transporter activity | 0.007651 | 0.007121 |
| **GSEA-BP** | GO:0001704 | formation of primary germ layer | 3.93E-05 | 3.61E-05 |
| **GSEA-BP** | GO:0001706 | endoderm formation | 3.93E-05 | 3.61E-05 |
| **GSEA-BP** | GO:0035987 | endodermal cell differentiation | 0.000437 | 0.000401 |
| **GSEA-BP** | GO:0048568 | embryonic organ development | 0.0007 | 0.000643 |
| **GSEA-BP** | GO:0042663 | regulation of endodermal cell fate specification | 0.003614 | 0.003319 |
| **GSEA-BP** | GO:0001707 | mesoderm formation | 0.007485 | 0.006876 |
| **GSEA-BP** | GO:0007492 | endoderm development | 0.007485 | 0.006876 |
| **GSEA-BP** | GO:0001711 | endodermal cell fate commitment | 0.007641 | 0.007018 |
| **GSEA-BP** | GO:0048562 | embryonic organ morphogenesis | 0.007641 | 0.007018 |
| **GSEA-BP** | GO:1904888 | cranial skeletal system development | 0.010401 | 0.009554 |
| **GSEA-BP** | GO:0048264 | determination of ventral identity | 0.01053 | 0.009672 |
| **GSEA-BP** | GO:0003002 | regionalization | 0.01053 | 0.009672 |
| **GSEA-BP** | GO:0045995 | regulation of embryonic development | 0.010555 | 0.009696 |
| **GSEA-BP** | GO:0001714 | endodermal cell fate specification | 0.013388 | 0.012298 |
| **GSEA-BP** | GO:0007389 | pattern specification process | 0.013488 | 0.01239 |
| **GSEA-BP** | GO:1903224 | regulation of endodermal cell differentiation | 0.014774 | 0.013571 |
| **GSEA-BP** | GO:0045165 | cell fate commitment | 0.014774 | 0.013571 |

| | | | | |
|---|---|---|---|---|
| **GSEA-BP** | GO:0048332 | mesoderm morphogenesis | 0.014811 | 0.013605 |
| **GSEA-BP** | GO:0060795 | cell fate commitment involved in formation of primary germ layer | 0.015018 | 0.013795 |
| **GSEA-BP** | GO:0042659 | regulation of cell fate specification | 0.015018 | 0.013795 |
| **GSEA-BP** | GO:0048729 | tissue morphogenesis | 0.015018 | 0.013795 |
| **GSEA-BP** | GO:0071600 | otic vesicle morphogenesis | 0.015658 | 0.014383 |
| **GSEA-BP** | GO:0001708 | cell fate specification | 0.023376 | 0.021472 |
| **GSEA-BP** | GO:0030916 | otic vesicle formation | 0.024311 | 0.022331 |
| **GSEA-BP** | GO:0007507 | heart development | 0.024311 | 0.022331 |
| **GSEA-BP** | GO:0035239 | tube morphogenesis | 0.026206 | 0.024072 |
| **GSEA-BP** | GO:0060788 | ectodermal placode formation | 0.039025 | 0.035846 |
| **GSEA-BP** | GO:0090092 | regulation of transmembrane receptor protein serine/threonine kinase signalling pathway | 0.039025 | 0.035846 |
| **GSEA-BP** | GO:0048701 | embryonic cranial skeleton morphogenesis | 0.039025 | 0.035846 |
| **GSEA-BP** | GO:0009952 | anterior/posterior pattern specification | 0.039025 | 0.035846 |
| **GSEA-BP** | GO:0060562 | epithelial tube morphogenesis | 0.039025 | 0.035846 |
| **GSEA-BP** | GO:0007369 | gastrulation | 0.039025 | 0.035846 |
| **GSEA-BP** | GO:0040011 | locomotion | 0.039025 | 0.035846 |
| **GSEA-BP** | GO:0007167 | enzyme-linked receptor protein signalling pathway | 0.039025 | 0.035846 |
| **GSEA-BP** | GO:0009719 | response to endogenous stimulus | 0.039025 | 0.035846 |
| **GSEA-BP** | GO:0002009 | morphogenesis of an epithelium | 0.039025 | 0.035846 |
| **GSEA-BP** | GO:0048870 | cell motility | 0.039025 | 0.035846 |
| **GSEA-BP** | GO:0016477 | cell migration | 0.039025 | 0.035846 |

| | | | | |
|---|---|---|---|---|
| **GSEA-BP** | GO:0043049 | otic placode formation | 0.040299 | 0.037017 |
| **GSEA-BP** | GO:0009953 | dorsal/ventral pattern formation | 0.040299 | 0.037017 |
| **GSEA-BP** | GO:0010453 | regulation of cell fate commitment | 0.042217 | 0.038779 |
| **GSEA-BP** | GO:0030513 | positive regulation of BMP signalling pathway | 0.04416 | 0.040564 |
| **GSEA-BP** | GO:0009798 | axis specification | 0.04416 | 0.040564 |
| **GSEA-CC** | GO:0005840 | ribosome | 0.006141 | 0.005871 |
| **GSEA-CC** | GO:0000313 | organellar ribosome | 0.009421 | 0.009008 |
| **GSEA-CC** | GO:0005761 | mitochondrial ribosome | 0.009421 | 0.009008 |
| **GSEA-MF** | GO:0038023 | signalling receptor activity | 5.33E-08 | 5.07E-08 |
| **GSEA-MF** | GO:0060089 | molecular transducer activity | 5.33E-08 | 5.07E-08 |
| **GSEA-MF** | GO:0004888 | transmembrane signalling receptor activity | 8.95E-05 | 8.51E-05 |
| **GSEA-MF** | GO:0005212 | structural constituent of eye lens | 0.005549 | 0.005271 |
| **GSEA-MF** | GO:0008146 | sulfotransferase activity | 0.049956 | 0.047457 |
| **GSEA-MF** | GO:0004930 | G protein-coupled receptor activity | 0.049956 | 0.047457 |
| **KEGG** | dre00190 | Oxidative phosphorylation | 0.000905 | 0.000871 |
| **KEGG** | dre04260 | Cardiac muscle contraction | 0.028293 | 0.027229 |
| **KEGG** | dre00500 | Starch and sucrose metabolism | 0.046695 | 0.008192 |
| **KEGG** | dre00480 | Glutathione metabolism | 0.046695 | 0.008192 |
| **KEGG** | dre00982 | Drug metabolism - cytochrome P450 | 0.046695 | 0.008192 |
| **KEGG** | dre00980 | Metabolism of xenobiotics by cytochrome P450 | 0.046695 | 0.008192 |

**Supplementary Table 3 | Functional enrichment analysis for the sphere stage (MZ*rlig1* vs. WT).** Listed are significantly (padj < 0.05) enriched terms from Gene Ontology (GO) subdivided into biological process (BP), cellular component (CC), and molecular function (MF), gene set enrichment analysis (GSEA) terms, and KEGG pathways.

| Term | ID | Description | padj | q-value |
|---|---|---|---|---|
| **GO-BP** | GO:0043065 | positive regulation of apoptotic process | 0.000794 | 0.000713 |
| **GO-BP** | GO:0043068 | positive regulation of programmed cell death | 0.000794 | 0.000713 |
| **GO-BP** | GO:0006091 | generation of precursor metabolites and energy | 0.001911 | 0.001715 |
| **GO-BP** | GO:0022904 | respiratory electron transport chain | 0.001957 | 0.001757 |
| **GO-BP** | GO:0042773 | ATP synthesis coupled electron transport | 0.001957 | 0.001757 |
| **GO-BP** | GO:0006119 | oxidative phosphorylation | 0.001957 | 0.001757 |
| **GO-BP** | GO:0022900 | electron transport chain | 0.002296 | 0.002061 |
| **GO-BP** | GO:0009060 | aerobic respiration | 0.002451 | 0.0022 |
| **GO-BP** | GO:0045333 | cellular respiration | 0.003439 | 0.003087 |
| **GO-BP** | GO:2000045 | regulation of G1/S transition of mitotic cell cycle | 0.004458 | 0.004001 |
| **GO-BP** | GO:0006120 | mitochondrial electron transport, NADH to ubiquinone | 0.007017 | 0.006299 |
| **GO-BP** | GO:1902806 | regulation of cell cycle G1/S phase transition | 0.007017 | 0.006299 |
| **GO-BP** | GO:0015980 | energy derivation by oxidation of organic compounds | 0.009395 | 0.008433 |
| **GO-BP** | GO:0051346 | negative regulation of hydrolase activity | 0.010857 | 0.009746 |
| **GO-BP** | GO:0007268 | chemical synaptic transmission | 0.010857 | 0.009746 |
| **GO-BP** | GO:0098916 | anterograde trans-synaptic signalling | 0.010857 | 0.009746 |
| **GO-BP** | GO:0042981 | regulation of apoptotic process | 0.010857 | 0.009746 |
| **GO-BP** | GO:0099537 | trans-synaptic signalling | 0.010857 | 0.009746 |
| **GO-BP** | GO:0019646 | aerobic electron transport chain | 0.011652 | 0.010459 |
| **GO-BP** | GO:0043067 | regulation of programmed cell death | 0.013368 | 0.011999 |
| **GO-BP** | GO:0042775 | mitochondrial ATP synthesis coupled electron transport | 0.013727 | 0.012321 |

| | | | | |
|---|---|---|---|---|
| **GO-BP** | GO:0051240 | positive regulation of multicellular organismal process | 0.013727 | 0.012321 |
| **GO-BP** | GO:0052547 | regulation of peptidase activity | 0.013727 | 0.012321 |
| **GO-BP** | GO:0099536 | synaptic signalling | 0.013728 | 0.012322 |
| **GO-BP** | GO:0007267 | cell-cell signalling | 0.016995 | 0.015255 |
| **GO-BP** | GO:0006915 | apoptotic process | 0.016995 | 0.015255 |
| **GO-BP** | GO:0043086 | negative regulation of catalytic activity | 0.016995 | 0.015255 |
| **GO-BP** | GO:0097190 | apoptotic signalling pathway | 0.019334 | 0.017355 |
| **GO-BP** | GO:1901616 | organic hydroxy compound catabolic process | 0.020377 | 0.018291 |
| **GO-BP** | GO:0008219 | cell death | 0.022881 | 0.020538 |
| **GO-BP** | GO:0012501 | programmed cell death | 0.022881 | 0.020538 |
| **GO-BP** | GO:0071391 | cellular response to estrogen stimulus | 0.026968 | 0.024207 |
| **GO-BP** | GO:0044092 | negative regulation of molecular function | 0.030928 | 0.027761 |
| **GO-BP** | GO:0043627 | response to estrogen | 0.035696 | 0.032041 |
| **GO-BP** | GO:1903037 | regulation of leukocyte cell-cell adhesion | 0.041872 | 0.037585 |
| **GO-BP** | GO:0035987 | endodermal cell differentiation | 0.049921 | 0.04481 |
| **GO-BP** | GO:0007159 | leukocyte cell-cell adhesion | 0.049921 | 0.04481 |
| **GO-BP** | GO:0010212 | response to ionizing radiation | 0.049921 | 0.04481 |
| **GO-BP** | GO:0010466 | negative regulation of peptidase activity | 0.049921 | 0.04481 |
| **GO-BP** | GO:0010951 | negative regulation of endopeptidase activity | 0.049921 | 0.04481 |
| **GO-BP** | GO:0042663 | regulation of endodermal cell fate specification | 0.049921 | 0.04481 |
| **GO-BP** | GO:0050870 | positive regulation of T cell activation | 0.049921 | 0.04481 |
| **GO-BP** | GO:1903039 | positive regulation of leukocyte cell-cell adhesion | 0.049921 | 0.04481 |
| **GO-BP** | GO:1905952 | regulation of lipid localization | 0.049921 | 0.04481 |
| **GO-CC** | GO:0005615 | extracellular space | 1.52E-07 | 1.37E-07 |
| **GO-CC** | GO:0005576 | extracellular region | 5.74E-06 | 5.16E-06 |
| **GO-CC** | GO:0070469 | respirasome | 7.74E-05 | 6.96E-05 |
| **GO-CC** | GO:0022626 | cytosolic ribosome | 0.000261 | 0.000235 |
| **GO-CC** | GO:0098803 | respiratory chain complex | 0.00059 | 0.000531 |
| **GO-CC** | GO:0022627 | cytosolic small ribosomal subunit | 0.00059 | 0.000531 |

| | | | | |
|---|---|---|---|---|
| **GO-CC** | GO:1990351 | transporter complex | 0.001728 | 0.001555 |
| **GO-CC** | GO:0005746 | mitochondrial respirasome | 0.001728 | 0.001555 |
| **GO-CC** | GO:1990204 | oxidoreductase complex | 0.003292 | 0.002962 |
| **GO-CC** | GO:1902495 | transmembrane transporter complex | 0.003292 | 0.002962 |
| **GO-CC** | GO:0005882 | intermediate filament | 0.005417 | 0.004874 |
| **GO-CC** | GO:0005747 | mitochondrial respiratory chain complex I | 0.005417 | 0.004874 |
| **GO-CC** | GO:0030964 | NADH dehydrogenase complex | 0.005417 | 0.004874 |
| **GO-CC** | GO:0045271 | respiratory chain complex I | 0.005417 | 0.004874 |
| **GO-CC** | GO:0098800 | inner mitochondrial membrane protein complex | 0.006986 | 0.006286 |
| **GO-CC** | GO:0045111 | intermediate filament cytoskeleton | 0.006986 | 0.006286 |
| **GO-CC** | GO:0015935 | small ribosomal subunit | 0.01208 | 0.01087 |
| **GO-CC** | GO:0044391 | ribosomal subunit | 0.016603 | 0.01494 |
| **GO-CC** | GO:0005840 | ribosome | 0.045196 | 0.040668 |
| **GO-MF** | GO:0015453 | oxidoreduction-driven active transmembrane transporter activity | 0.000239 | 0.000215 |
| **GO-MF** | GO:0009055 | electron transfer activity | 0.000239 | 0.000215 |
| **GO-MF** | GO:0003954 | NADH dehydrogenase activity | 0.001648 | 0.001481 |
| **GO-MF** | GO:0008137 | NADH dehydrogenase (ubiquinone) activity | 0.002715 | 0.00244 |
| **GO-MF** | GO:0050136 | NADH dehydrogenase (quinone) activity | 0.002715 | 0.00244 |
| **GO-MF** | GO:0048018 | receptor ligand activity | 0.002715 | 0.00244 |
| **GO-MF** | GO:0030546 | signalling receptor activator activity | 0.002954 | 0.002656 |
| **GO-MF** | GO:0003955 | NAD(P)H dehydrogenase (quinone) activity | 0.003035 | 0.002728 |
| **GO-MF** | GO:0030545 | signalling receptor regulator activity | 0.003057 | 0.002747 |
| **GO-MF** | GO:0005102 | signalling receptor binding | 0.003057 | 0.002747 |
| **GO-MF** | GO:0005179 | hormone activity | 0.003057 | 0.002747 |
| **GO-MF** | GO:0016655 | oxidoreductase activity, acting on NAD(P)H, quinone or similar compound as acceptor | 0.003057 | 0.002747 |
| **GO-MF** | GO:0004857 | enzyme inhibitor activity | 0.004088 | 0.003675 |
| **GO-MF** | GO:0140678 | molecular function inhibitor activity | 0.004162 | 0.003741 |
| **GO-MF** | GO:0061134 | peptidase regulator activity | 0.01019 | 0.009159 |

| | | | | |
|---|---|---|---|---|
| **GO-MF** | GO:0016651 | oxidoreductase activity, acting on NAD(P)H | 0.018103 | 0.016271 |
| **GO-MF** | GO:0003735 | structural constituent of ribosome | 0.022525 | 0.020246 |
| **GO-MF** | GO:0015399 | primary active transmembrane transporter activity | 0.030928 | 0.027799 |
| **GO-MF** | GO:0061135 | endopeptidase regulator activity | 0.046208 | 0.041533 |
| **GSEA-BP** | GO:0042254 | ribosome biogenesis | 0.016387 | 0.015619 |
| **GSEA-CC** | GO:0022626 | cytosolic ribosome | 1.47E-08 | 1.33E-08 |
| **GSEA-CC** | GO:0044391 | ribosomal subunit | 1.47E-08 | 1.33E-08 |
| **GSEA-CC** | GO:0005840 | ribosome | 1.47E-08 | 1.33E-08 |
| **GSEA-CC** | GO:0022625 | cytosolic large ribosomal subunit | 1.06E-06 | 9.59E-07 |
| **GSEA-CC** | GO:0005576 | extracellular region | 6.06E-06 | 5.47E-06 |
| **GSEA-CC** | GO:0022627 | cytosolic small ribosomal subunit | 1.99E-05 | 1.8E-05 |
| **GSEA-CC** | GO:0015934 | large ribosomal subunit | 2.6E-05 | 2.35E-05 |
| **GSEA-CC** | GO:0005615 | extracellular space | 5.34E-05 | 4.82E-05 |
| **GSEA-CC** | GO:0005882 | intermediate filament | 0.000278 | 0.000251 |
| **GSEA-CC** | GO:0045111 | intermediate filament cytoskeleton | 0.000917 | 0.000828 |
| **GSEA-CC** | GO:0015935 | small ribosomal subunit | 0.001019 | 0.00092 |
| **GSEA-CC** | GO:0015630 | microtubule cytoskeleton | 0.001713 | 0.001546 |
| **GSEA-CC** | GO:0030312 | external encapsulating structure | 0.004447 | 0.004014 |
| **GSEA-CC** | GO:0031012 | extracellular matrix | 0.004447 | 0.004014 |
| **GSEA-CC** | GO:0062023 | collagen-containing extracellular matrix | 0.01017 | 0.00918 |
| **GSEA-CC** | GO:0070469 | respirasome | 0.018944 | 0.017099 |
| **GSEA-CC** | GO:0005815 | microtubule organizing center | 0.037673 | 0.034004 |
| **GSEA-CC** | GO:0005730 | nucleolus | 0.038533 | 0.03478 |
| **GSEA-CC** | GO:1990204 | oxidoreductase complex | 0.046997 | 0.04242 |
| **GSEA-MF** | GO:0003735 | structural constituent of ribosome | 3.49E-08 | 3.25E-08 |
| **GSEA-MF** | GO:0005198 | structural molecule activity | 3.49E-08 | 3.25E-08 |
| **GSEA-MF** | GO:0030695 | GTPase regulator activity | 0.000721 | 0.000673 |
| **GSEA-MF** | GO:0060589 | nucleoside-triphosphatase regulator activity | 0.000721 | 0.000673 |
| **GSEA-MF** | GO:0030165 | PDZ domain binding | 0.008757 | 0.008173 |
| **GSEA-MF** | GO:0004672 | protein kinase activity | 0.010105 | 0.009431 |
| **GSEA-MF** | GO:0009055 | electron transfer activity | 0.01608 | 0.015007 |
| **GSEA-MF** | GO:0003954 | NADH dehydrogenase activity | 0.019065 | 0.017794 |

| | | | | |
|---|---|---|---|---|
| **GSEA-MF** | GO:0005096 | GTPase activator activity | 0.031941 | 0.029812 |
| **GSEA-MF** | GO:0015453 | oxidoreduction-driven active transmembrane transporter activity | 0.038277 | 0.035725 |
| **GSEA-MF** | GO:0019843 | rRNA binding | 0.039347 | 0.036723 |
| **GSEA-MF** | GO:0031492 | nucleosomal DNA binding | 0.039347 | 0.036723 |
| **GSEA-MF** | GO:0016620 | oxidoreductase activity, acting on the aldehyde or oxo group of donors, NAD or NADP as acceptor | 0.039347 | 0.036723 |
| **GSEA-MF** | GO:0008137 | NADH dehydrogenase (ubiquinone) activity | 0.039347 | 0.036723 |
| **GSEA-MF** | GO:0050136 | NADH dehydrogenase (quinone) activity | 0.039347 | 0.036723 |
| **GSEA-MF** | GO:0005085 | guanyl-nucleotide exchange factor activity | 0.039347 | 0.036723 |
| **GSEA-MF** | GO:0008017 | microtubule binding | 0.039347 | 0.036723 |
| **GSEA-MF** | GO:0004721 | phosphoprotein phosphatase activity | 0.039347 | 0.036723 |
| **GSEA-MF** | GO:0042578 | phosphoric ester hydrolase activity | 0.039347 | 0.036723 |
| **GSEA-MF** | GO:0016791 | phosphatase activity | 0.039347 | 0.036723 |
| **GSEA-MF** | GO:0005230 | extracellular ligand-gated monoatomic ion channel activity | 0.042391 | 0.039564 |
| **KEGG** | dre00190 | Oxidative phosphorylation | 0.000116 | 0.000106 |
| **KEGG** | dre00051 | Fructose and mannose metabolism | 0.005258 | 0.004837 |
| **KEGG** | dre00030 | Pentose phosphate pathway | 0.009528 | 0.008765 |
| **KEGG** | dre04115 | p53 signalling pathway | 0.025545 | 0.023498 |

**Supplementary Table 4 | Functional enrichment analysis for the shield stage (MZ*rlig1* vs. WT).** Listed are significantly (padj < 0.05) enriched terms from Gene Ontology (GO) subdivided into biological process (BP), cellular component (CC), and molecular function (MF), gene set enrichment analysis (GSEA) terms, and KEGG pathways.

| Term | ID | Description | padj | q-value |
|---|---|---|---|---|
| **GO-BP** | GO:0048568 | embryonic organ development | 8.54E-06 | 7.85E-06 |
| **GO-BP** | GO:0048839 | inner ear development | 0.000298 | 0.000274 |
| **GO-BP** | GO:0043583 | ear development | 0.000298 | 0.000274 |
| **GO-BP** | GO:0048562 | embryonic organ morphogenesis | 0.00192 | 0.001766 |
| **GO-BP** | GO:0090596 | sensory organ morphogenesis | 0.005504 | 0.005063 |
| **GO-BP** | GO:0031076 | embryonic camera-type eye development | 0.00609 | 0.005602 |
| **GO-BP** | GO:0042472 | inner ear morphogenesis | 0.011177 | 0.010282 |
| **GO-BP** | GO:0060322 | head development | 0.011177 | 0.010282 |
| **GO-BP** | GO:0042471 | ear morphogenesis | 0.011177 | 0.010282 |
| **GO-BP** | GO:0043009 | chordate embryonic development | 0.011177 | 0.010282 |
| **GO-BP** | GO:0048729 | tissue morphogenesis | 0.011177 | 0.010282 |
| **GO-BP** | GO:0009792 | embryo development ending in birth or egg hatching | 0.011177 | 0.010282 |
| **GO-BP** | GO:0007420 | brain development | 0.01144 | 0.010523 |
| **GO-BP** | GO:0048880 | sensory system development | 0.011939 | 0.010982 |
| **GO-BP** | GO:0035148 | tube formation | 0.013125 | 0.012073 |
| **GO-BP** | GO:0072175 | epithelial tube formation | 0.043146 | 0.039688 |
| **GO-BP** | GO:0001654 | eye development | 0.046653 | 0.042914 |
| **GO-BP** | GO:0150063 | visual system development | 0.046653 | 0.042914 |
| **GO-BP** | GO:0048762 | mesenchymal cell differentiation | 0.046831 | 0.043078 |
| **GO-CC** | GO:0005930 | axoneme | 0.043816 | 0.043409 |
| **GO-CC** | GO:0097014 | ciliary plasm | 0.043816 | 0.043409 |
| **GO-CC** | GO:0032838 | plasma membrane bounded cell projection cytoplasm | 0.043816 | 0.043409 |
| **GSEA-BP** | GO:0060485 | mesenchyme development | 0.008653 | 0.008444 |
| **GSEA-BP** | GO:0048863 | stem cell differentiation | 0.030177 | 0.029451 |
| **GSEA-BP** | GO:0007507 | heart development | 0.039889 | 0.038929 |
| **GSEA-BP** | GO:0048568 | embryonic organ development | 0.042194 | 0.041178 |
| **GSEA-BP** | GO:0035725 | sodium ion transmembrane transport | 0.047 | 0.045869 |
| **GSEA-BP** | GO:0014032 | neural crest cell development | 0.047 | 0.045869 |
| **GSEA-BP** | GO:0048864 | stem cell development | 0.047 | 0.045869 |

| | | | | |
|---|---|---|---|---|
| **GSEA-BP** | GO:0014033 | neural crest cell differentiation | 0.047 | 0.045869 |
| **GSEA-BP** | GO:0048762 | mesenchymal cell differentiation | 0.047 | 0.045869 |
| **GSEA-BP** | GO:0060322 | head development | 0.047 | 0.045869 |
| **GSEA-BP** | GO:0043009 | chordate embryonic development | 0.047 | 0.045869 |
| **GSEA-BP** | GO:0009792 | embryo development ending in birth or egg hatching | 0.047 | 0.045869 |
| **GSEA-CC** | GO:0000786 | nucleosome | 0.000739 | 0.000707 |
| **GSEA-CC** | GO:0043005 | neuron projection | 0.017256 | 0.016509 |
| **GSEA-CC** | GO:0022626 | cytosolic ribosome | 0.018949 | 0.018129 |
| **GSEA-CC** | GO:0005929 | cilium | 0.021316 | 0.020394 |
| **GSEA-MF** | GO:0030527 | structural constituent of chromatin | 0.007611 | 0.007215 |
| **KEGG** | dre04382 | Cornified envelope formation | 0.003301 | 0.003145 |

**Supplementary Table 5 | Functional enrichment analysis for the bud stage (MZ*rlig1* vs. WT).** Listed are significantly (padj < 0.05) enriched terms from Gene Ontology (GO) subdivided into biological process (BP), cellular component (CC), and molecular function (MF), gene set enrichment analysis (GSEA) terms, and KEGG pathways.

| Term | ID | Description | padj | q-value |
|---|---|---|---|---|
| **GO-BP** | GO:0035249 | synaptic transmission, glutamatergic | 0.0364 | 0.034338 |
| **GO-BP** | GO:0060041 | retina development in camera-type eye | 0.0364 | 0.034338 |
| **GO-BP** | GO:0042063 | gliogenesis | 0.03883 | 0.03663 |
| **GO-BP** | GO:0090596 | sensory organ morphogenesis | 0.03883 | 0.03663 |
| **GO-BP** | GO:0043010 | camera-type eye development | 0.03883 | 0.03663 |
| **GO-BP** | GO:0050935 | iridophore differentiation | 0.03883 | 0.03663 |
| **GO-BP** | GO:0010001 | glial cell differentiation | 0.048077 | 0.045353 |
| **GO-BP** | GO:0048593 | camera-type eye morphogenesis | 0.048077 | 0.045353 |
| **GO-CC** | GO:0098839 | postsynaptic density membrane | 0.00612 | 0.00544 |
| **GO-CC** | GO:0099634 | postsynaptic specialization membrane | 0.00612 | 0.00544 |
| **GO-CC** | GO:0014069 | postsynaptic density | 0.00612 | 0.00544 |
| **GO-CC** | GO:0032279 | asymmetric synapse | 0.00612 | 0.00544 |
| **GO-CC** | GO:0098984 | neuron to neuron synapse | 0.00612 | 0.00544 |
| **GO-CC** | GO:0099572 | postsynaptic specialization | 0.007017 | 0.006238 |
| **GO-CC** | GO:0008328 | ionotropic glutamate receptor complex | 0.021738 | 0.019322 |
| **GO-CC** | GO:0098878 | neurotransmitter receptor complex | 0.021738 | 0.019322 |
| **GO-CC** | GO:0030312 | external encapsulating structure | 0.039455 | 0.035071 |
| **GO-CC** | GO:0031012 | extracellular matrix | 0.039455 | 0.035071 |
| **GO-MF** | GO:0008066 | glutamate receptor activity | 0.001014 | 0.000923 |
| **GO-MF** | GO:0004970 | ionotropic glutamate receptor activity | 0.003815 | 0.003471 |
| **GO-MF** | GO:0099529 | neurotransmitter receptor activity involved in regulation of postsynaptic membrane potential | 0.013556 | 0.012334 |
| **GO-MF** | GO:1904315 | transmitter-gated monoatomic ion channel activity involved in regulation of postsynaptic membrane potential | 0.013556 | 0.012334 |

| | | | | |
|---|---|---|---|---|
| **GO-MF** | GO:0022824 | transmitter-gated monoatomic ion channel activity | 0.020209 | 0.018387 |
| **GO-MF** | GO:0022835 | transmitter-gated channel activity | 0.020209 | 0.018387 |
| **GO-MF** | GO:0098960 | postsynaptic neurotransmitter receptor activity | 0.020209 | 0.018387 |
| **GO-MF** | GO:0005230 | extracellular ligand-gated monoatomic ion channel activity | 0.0315 | 0.02866 |
| **GO-MF** | GO:0004888 | transmembrane signaling receptor activity | 0.034736 | 0.031604 |
| **GO-MF** | GO:0015276 | ligand-gated monoatomic ion channel activity | 0.034736 | 0.031604 |
| **GO-MF** | GO:0022834 | ligand-gated channel activity | 0.034736 | 0.031604 |
| **GO-MF** | GO:0005272 | sodium channel activity | 0.035084 | 0.03192 |
| **GSEA-BP** | GO:2000241 | regulation of reproductive process | 0.000412 | 0.000382 |
| **GSEA-BP** | GO:2000243 | positive regulation of reproductive process | 0.01186 | 0.010998 |
| **GSEA-BP** | GO:0007339 | binding of sperm to zona pellucida | 0.01186 | 0.010998 |
| **GSEA-BP** | GO:0035036 | sperm-egg recognition | 0.01186 | 0.010998 |
| **GSEA-BP** | GO:0060348 | bone development | 0.01186 | 0.010998 |
| **GSEA-BP** | GO:0048709 | oligodendrocyte differentiation | 0.01186 | 0.010998 |
| **GSEA-BP** | GO:0014706 | striated muscle tissue development | 0.01186 | 0.010998 |
| **GSEA-BP** | GO:0048738 | cardiac muscle tissue development | 0.01186 | 0.010998 |
| **GSEA-BP** | GO:0010001 | glial cell differentiation | 0.01186 | 0.010998 |
| **GSEA-BP** | GO:0042063 | gliogenesis | 0.01186 | 0.010998 |
| **GSEA-BP** | GO:0009988 | cell-cell recognition | 0.013991 | 0.012975 |
| **GSEA-BP** | GO:0035265 | organ growth | 0.013991 | 0.012975 |
| **GSEA-BP** | GO:0006334 | nucleosome assembly | 0.017657 | 0.016374 |
| **GSEA-BP** | GO:0021782 | glial cell development | 0.017657 | 0.016374 |
| **GSEA-BP** | GO:0045229 | external encapsulating structure organization | 0.017657 | 0.016374 |
| **GSEA-BP** | GO:0022613 | ribonucleoprotein complex biogenesis | 0.017657 | 0.016374 |
| **GSEA-BP** | GO:0034728 | nucleosome organization | 0.018797 | 0.017431 |
| **GSEA-BP** | GO:0035082 | axoneme assembly | 0.018797 | 0.017431 |
| **GSEA-BP** | GO:0065004 | protein-DNA complex assembly | 0.018797 | 0.017431 |
| **GSEA-BP** | GO:0030198 | extracellular matrix organization | 0.018797 | 0.017431 |

| | | | | |
|---|---|---|---|---|
| **GSEA-BP** | GO:0043062 | extracellular structure organization | 0.018797 | 0.017431 |
| **GSEA-BP** | GO:0060537 | muscle tissue development | 0.018797 | 0.017431 |
| **GSEA-BP** | GO:0042254 | ribosome biogenesis | 0.018797 | 0.017431 |
| **GSEA-BP** | GO:0061061 | muscle structure development | 0.019217 | 0.017821 |
| **GSEA-BP** | GO:0061564 | axon development | 0.019217 | 0.017821 |
| **GSEA-BP** | GO:0001501 | skeletal system development | 0.021926 | 0.020333 |
| **GSEA-BP** | GO:0007411 | axon guidance | 0.022277 | 0.020659 |
| **GSEA-BP** | GO:0097485 | neuron projection guidance | 0.022277 | 0.020659 |
| **GSEA-BP** | GO:0048589 | developmental growth | 0.031804 | 0.029494 |
| **GSEA-BP** | GO:0032474 | otolith morphogenesis | 0.037607 | 0.034875 |
| **GSEA-BP** | GO:0001654 | eye development | 0.037607 | 0.034875 |
| **GSEA-BP** | GO:0150063 | visual system development | 0.037607 | 0.034875 |
| **GSEA-BP** | GO:0022414 | reproductive process | 0.037655 | 0.03492 |
| **GSEA-BP** | GO:0040007 | growth | 0.039501 | 0.036632 |
| **GSEA-BP** | GO:0055017 | cardiac muscle tissue growth | 0.047986 | 0.044501 |
| **GSEA-BP** | GO:0098609 | cell-cell adhesion | 0.048356 | 0.044843 |
| **GSEA-BP** | GO:0000003 | reproduction | 0.048356 | 0.044843 |
| **GSEA-BP** | GO:0048562 | embryonic organ morphogenesis | 0.048356 | 0.044843 |
| **GSEA-BP** | GO:0043010 | camera-type eye development | 0.048356 | 0.044843 |
| **GSEA-BP** | GO:0042692 | muscle cell differentiation | 0.048772 | 0.045229 |
| **GSEA-CC** | GO:0022626 | cytosolic ribosome | 4.84E-08 | 4.26E-08 |
| **GSEA-CC** | GO:0044391 | ribosomal subunit | 1.87E-07 | 1.65E-07 |
| **GSEA-CC** | GO:0005840 | ribosome | 2.82E-07 | 2.48E-07 |
| **GSEA-CC** | GO:0000786 | nucleosome | 2.22E-06 | 1.95E-06 |
| **GSEA-CC** | GO:0022625 | cytosolic large ribosomal subunit | 0.000282 | 0.000248 |
| **GSEA-CC** | GO:0030312 | external encapsulating structure | 0.000282 | 0.000248 |
| **GSEA-CC** | GO:0031012 | extracellular matrix | 0.000282 | 0.000248 |
| **GSEA-CC** | GO:0062023 | collagen-containing extracellular matrix | 0.000816 | 0.000719 |
| **GSEA-CC** | GO:0022627 | cytosolic small ribosomal subunit | 0.000854 | 0.000753 |
| **GSEA-CC** | GO:0015935 | small ribosomal subunit | 0.000913 | 0.000804 |
| **GSEA-CC** | GO:0015934 | large ribosomal subunit | 0.001274 | 0.001122 |
| **GSEA-CC** | GO:0098797 | plasma membrane protein complex | 0.011499 | 0.010128 |
| **GSEA-CC** | GO:0005604 | basement membrane | 0.017337 | 0.01527 |
| **GSEA-CC** | GO:0043235 | receptor complex | 0.019525 | 0.017198 |
| **GSEA-CC** | GO:0035805 | egg coat | 0.021102 | 0.018587 |
| **GSEA-CC** | GO:0005581 | collagen trimer | 0.021102 | 0.018587 |
| **GSEA-CC** | GO:0043005 | neuron projection | 0.021102 | 0.018587 |

| | | | | |
|---|---|---|---|---|
| **GSEA-CC** | GO:0034704 | calcium channel complex | 0.021246 | 0.018714 |
| **GSEA-CC** | GO:0030672 | synaptic vesicle membrane | 0.024088 | 0.021217 |
| **GSEA-CC** | GO:0099501 | exocytic vesicle membrane | 0.024088 | 0.021217 |
| **GSEA-CC** | GO:0031514 | motile cilium | 0.026283 | 0.02315 |
| **GSEA-CC** | GO:0005891 | voltage-gated calcium channel complex | 0.02947 | 0.025957 |
| **GSEA-CC** | GO:0000502 | proteasome complex | 0.029567 | 0.026043 |
| **GSEA-CC** | GO:0099081 | supramolecular polymer | 0.029567 | 0.026043 |
| **GSEA-CC** | GO:0099512 | supramolecular fiber | 0.029567 | 0.026043 |
| **GSEA-MF** | GO:0003735 | structural constituent of ribosome | 8.29E-08 | 7.81E-08 |
| **GSEA-MF** | GO:0030527 | structural constituent of chromatin | 8.72E-07 | 8.22E-07 |
| **GSEA-MF** | GO:0004888 | transmembrane signalling receptor activity | 0.005003 | 0.004714 |
| **GSEA-MF** | GO:0022836 | gated channel activity | 0.018292 | 0.017234 |
| **GSEA-MF** | GO:0022839 | monoatomic ion gated channel activity | 0.018292 | 0.017234 |
| **GSEA-MF** | GO:0015267 | channel activity | 0.018292 | 0.017234 |
| **GSEA-MF** | GO:0022803 | passive transmembrane transporter activity | 0.018292 | 0.017234 |
| **GSEA-MF** | GO:0032190 | acrosin binding | 0.018756 | 0.017671 |
| **GSEA-MF** | GO:0098960 | postsynaptic neurotransmitter receptor activity | 0.018756 | 0.017671 |
| **GSEA-MF** | GO:0099529 | neurotransmitter receptor activity involved in regulation of postsynaptic membrane potential | 0.018756 | 0.017671 |
| **GSEA-MF** | GO:1904315 | transmitter-gated monoatomic ion channel activity involved in regulation of postsynaptic membrane potential | 0.018756 | 0.017671 |
| **GSEA-MF** | GO:0005216 | monoatomic ion channel activity | 0.018756 | 0.017671 |
| **GSEA-MF** | GO:0022824 | transmitter-gated monoatomic ion channel activity | 0.024794 | 0.02336 |
| **GSEA-MF** | GO:0022835 | transmitter-gated channel activity | 0.024794 | 0.02336 |
| **GSEA-MF** | GO:0030594 | neurotransmitter receptor activity | 0.024794 | 0.02336 |
| **GSEA-MF** | GO:0005261 | monoatomic cation channel activity | 0.030436 | 0.028675 |
| **GSEA-MF** | GO:0030020 | extracellular matrix structural constituent conferring tensile strength | 0.033152 | 0.031235 |
| **GSEA-MF** | GO:0004175 | endopeptidase activity | 0.045844 | 0.043193 |

| | | | | |
|---|---|---|---|---|
| **GSEA-MF** | GO:0035804 | structural constituent of egg coat | 0.046212 | 0.043539 |
| **GSEA-MF** | GO:0022848 | acetylcholine-gated monoatomic cation-selective channel activity | 0.046212 | 0.043539 |
| **GSEA-MF** | GO:0042562 | hormone binding | 0.047127 | 0.044402 |

**Supplementary Table 6 | Functional enrichment analysis for 1 dpf (MZ*rlig1* vs. WT).** Listed are significantly (padj < 0.05) enriched terms from Gene Ontology (GO) subdivided into biological process (BP), cellular component (CC), and molecular function (MF), gene set enrichment analysis (GSEA) terms, and KEGG pathways.

| Term | ID | Description | padj | q-value |
|---|---|---|---|---|
| **GO-BP** | GO:0002252 | immune effector process | 0.015924 | 0.007483 |
| **GO-BP** | GO:0031640 | killing of cells of another organism | 0.039139 | 0.018392 |
| **GO-BP** | GO:0141061 | disruption of cell in another organism | 0.039139 | 0.018392 |
| **GO-BP** | GO:0001906 | cell killing | 0.039139 | 0.018392 |
| **GO-BP** | GO:0141060 | disruption of anatomical structure in another organism | 0.039139 | 0.018392 |
| **GO-BP** | GO:0046653 | tetrahydrofolate metabolic process | 0.039139 | 0.018392 |
| **GO-BP** | GO:0009988 | cell-cell recognition | 0.039139 | 0.018392 |
| **GO-BP** | GO:0006955 | immune response | 0.039139 | 0.018392 |
| **GO-BP** | GO:0006760 | folic acid-containing compound metabolic process | 0.039139 | 0.018392 |
| **GO-BP** | GO:0006956 | complement activation | 0.041216 | 0.019369 |
| **GO-BP** | GO:0002449 | lymphocyte mediated immunity | 0.041216 | 0.019369 |
| **GO-BP** | GO:0042558 | pteridine-containing compound metabolic process | 0.041216 | 0.019369 |
| **GO-BP** | GO:0002460 | adaptive immune response based on somatic recombination of immune receptors built from immunoglobulin superfamily domains | 0.041216 | 0.019369 |
| **GO-BP** | GO:0048702 | embryonic neurocranium morphogenesis | 0.041216 | 0.019369 |
| **GO-BP** | GO:0006334 | nucleosome assembly | 0.041216 | 0.019369 |
| **GO-BP** | GO:0002443 | leukocyte mediated immunity | 0.041216 | 0.019369 |
| **GO-BP** | GO:0006959 | humoral immune response | 0.041216 | 0.019369 |
| **GO-BP** | GO:0034728 | nucleosome organization | 0.041216 | 0.019369 |
| **GO-BP** | GO:0006730 | one-carbon metabolic process | 0.041216 | 0.019369 |
| **GO-BP** | GO:0051607 | defense response to virus | 0.041216 | 0.019369 |
| **GO-BP** | GO:0140546 | defense response to symbiont | 0.041216 | 0.019369 |
| **GO-BP** | GO:0002250 | adaptive immune response | 0.041455 | 0.019481 |
| **GO-BP** | GO:0008037 | cell recognition | 0.045731 | 0.02149 |
| **GO-BP** | GO:0009615 | response to virus | 0.045731 | 0.02149 |
| **GO-CC** | GO:0046930 | pore complex | 5.5E-05 | 2.31E-05 |

| | | | | |
|---|---|---|---|---|
| **GO-CC** | GO:0098797 | plasma membrane protein complex | 0.017 | 0.007158 |
| **GO-CC** | GO:0000786 | nucleosome | 0.019803 | 0.008338 |
| **GO-CC** | GO:0005940 | septin ring | 0.009586 | NA |
| **GO-CC** | GO:0031105 | septin complex | 0.009586 | NA |
| **GO-CC** | GO:0032156 | septin cytoskeleton | 0.009586 | NA |
| **GO-CC** | GO:0032153 | cell division site | 0.018425 | NA |
| **GO-CC** | GO:0005938 | cell cortex | 0.037878 | NA |
| **GO-MF** | GO:0016874 | ligase activity | 0.030352 | 0.013452 |
| **GO-MF** | GO:0001671 | ATPase activator activity | 0.030352 | 0.013452 |
| **GO-MF** | GO:0016646 | oxidoreductase activity, acting on the CH-NH group of donors, NAD or NADP as acceptor | 0.030352 | 0.013452 |
| **GO-MF** | GO:0030527 | structural constituent of chromatin | 0.030352 | 0.013452 |
| **GO-MF** | GO:0016645 | oxidoreductase activity, acting on the CH-NH group of donors | 0.030352 | 0.013452 |
| **GO-MF** | GO:0022829 | wide pore channel activity | 0.030352 | 0.013452 |
| **GO-MF** | GO:0016814 | hydrolase activity, acting on carbon-nitrogen (but not peptide) bonds, in cyclic amidines | 0.030352 | 0.013452 |
| **GO-MF** | GO:0060590 | ATPase regulator activity | 0.030352 | 0.013452 |
| **GO-MF** | GO:0016879 | ligase activity, forming carbon-nitrogen bonds | 0.041125 | 0.018227 |
| **GO-MF** | GO:0051087 | protein-folding chaperone binding | 0.041125 | 0.018227 |
| **GO-MF** | GO:0031386 | protein tag activity | 0.018821 | 0.013208 |
| **GO-MF** | GO:0141047 | molecular tag activity | 0.018821 | 0.013208 |
| **GSEA-BP** | GO:0022613 | ribonucleoprotein complex biogenesis | 2.15E-05 | 2.04E-05 |
| **GSEA-BP** | GO:0043043 | peptide biosynthetic process | 2.96E-05 | 2.8E-05 |
| **GSEA-BP** | GO:0006412 | translation | 3.22E-05 | 3.05E-05 |
| **GSEA-BP** | GO:0006518 | peptide metabolic process | 0.000206 | 0.000195 |
| **GSEA-BP** | GO:0007599 | hemostasis | 0.000304 | 0.000287 |
| **GSEA-BP** | GO:0050878 | regulation of body fluid levels | 0.000452 | 0.000428 |
| **GSEA-BP** | GO:0007596 | blood coagulation | 0.000762 | 0.000722 |
| **GSEA-BP** | GO:0042254 | ribosome biogenesis | 0.001181 | 0.001118 |
| **GSEA-BP** | GO:0043604 | amide biosynthetic process | 0.002008 | 0.001902 |
| **GSEA-BP** | GO:0019646 | aerobic electron transport chain | 0.002171 | 0.002056 |
| **GSEA-BP** | GO:0050817 | coagulation | 0.002345 | 0.002221 |

| | | | | |
|---|---|---|---|---|
| **GSEA-BP** | GO:0042775 | mitochondrial ATP synthesis coupled electron transport | 0.003042 | 0.002881 |
| **GSEA-BP** | GO:0042773 | ATP synthesis coupled electron transport | 0.003377 | 0.003198 |
| **GSEA-BP** | GO:0006119 | oxidative phosphorylation | 0.003377 | 0.003198 |
| **GSEA-BP** | GO:0006935 | chemotaxis | 0.003377 | 0.003198 |
| **GSEA-BP** | GO:0042330 | taxis | 0.003377 | 0.003198 |
| **GSEA-BP** | GO:0003044 | regulation of systemic arterial blood pressure mediated by a chemical signal | 0.005714 | 0.005411 |
| **GSEA-BP** | GO:0009060 | aerobic respiration | 0.007245 | 0.006861 |
| **GSEA-BP** | GO:0022904 | respiratory electron transport chain | 0.007967 | 0.007545 |
| **GSEA-BP** | GO:0009144 | purine nucleoside triphosphate metabolic process | 0.007967 | 0.007545 |
| **GSEA-BP** | GO:0022900 | electron transport chain | 0.009044 | 0.008565 |
| **GSEA-BP** | GO:0045333 | cellular respiration | 0.009044 | 0.008565 |
| **GSEA-BP** | GO:0008610 | lipid biosynthetic process | 0.009274 | 0.008783 |
| **GSEA-BP** | GO:0030595 | leukocyte chemotaxis | 0.012437 | 0.011778 |
| **GSEA-BP** | GO:0042060 | wound healing | 0.013466 | 0.012753 |
| **GSEA-BP** | GO:0003073 | regulation of systemic arterial blood pressure | 0.014197 | 0.013445 |
| **GSEA-BP** | GO:0046467 | membrane lipid biosynthetic process | 0.014197 | 0.013445 |
| **GSEA-BP** | GO:0007005 | mitochondrion organization | 0.021089 | 0.019971 |
| **GSEA-BP** | GO:0006091 | generation of precursor metabolites and energy | 0.022502 | 0.021309 |
| **GSEA-BP** | GO:0009145 | purine nucleoside triphosphate biosynthetic process | 0.02458 | 0.023277 |
| **GSEA-BP** | GO:0009206 | purine ribonucleoside triphosphate biosynthetic process | 0.02458 | 0.023277 |
| **GSEA-BP** | GO:0046777 | protein autophosphorylation | 0.02458 | 0.023277 |
| **GSEA-BP** | GO:0015980 | energy derivation by oxidation of organic compounds | 0.02458 | 0.023277 |
| **GSEA-BP** | GO:0006364 | rRNA processing | 0.02458 | 0.023277 |
| **GSEA-BP** | GO:0050886 | endocrine process | 0.025004 | 0.023679 |
| **GSEA-BP** | GO:0060326 | cell chemotaxis | 0.025753 | 0.024388 |
| **GSEA-BP** | GO:0030148 | sphingolipid biosynthetic process | 0.025807 | 0.02444 |
| **GSEA-BP** | GO:0033108 | mitochondrial respiratory chain complex assembly | 0.02703 | 0.025598 |

| | | | | |
|---|---|---|---|---|
| **GSEA-BP** | GO:0019229 | regulation of vasoconstriction | 0.02703 | 0.025598 |
| **GSEA-BP** | GO:0016072 | rRNA metabolic process | 0.02703 | 0.025598 |
| **GSEA-BP** | GO:0070098 | chemokine-mediated signalling pathway | 0.03198 | 0.030286 |
| **GSEA-BP** | GO:1990868 | response to chemokine | 0.03319 | 0.031432 |
| **GSEA-BP** | GO:1990869 | cellular response to chemokine | 0.03319 | 0.031432 |
| **GSEA-BP** | GO:0009141 | nucleoside triphosphate metabolic process | 0.039644 | 0.037544 |
| **GSEA-BP** | GO:0071826 | protein-RNA complex organization | 0.047853 | 0.045318 |
| **GSEA-CC** | GO:0005840 | ribosome | 4.82E-09 | 4.14E-09 |
| **GSEA-CC** | GO:0098798 | mitochondrial protein-containing complex | 4.82E-09 | 4.14E-09 |
| **GSEA-CC** | GO:0044391 | ribosomal subunit | 4.82E-09 | 4.14E-09 |
| **GSEA-CC** | GO:0070469 | respirasome | 4.82E-09 | 4.14E-09 |
| **GSEA-CC** | GO:0098803 | respiratory chain complex | 4.82E-09 | 4.14E-09 |
| **GSEA-CC** | GO:0005746 | mitochondrial respirasome | 4.82E-09 | 4.14E-09 |
| **GSEA-CC** | GO:0098800 | inner mitochondrial membrane protein complex | 4.82E-09 | 4.14E-09 |
| **GSEA-CC** | GO:0005743 | mitochondrial inner membrane | 4.82E-09 | 4.14E-09 |
| **GSEA-CC** | GO:0019866 | organelle inner membrane | 4.82E-09 | 4.14E-09 |
| **GSEA-CC** | GO:1990904 | ribonucleoprotein complex | 4.82E-09 | 4.14E-09 |
| **GSEA-CC** | GO:0015934 | large ribosomal subunit | 1.03E-07 | 8.88E-08 |
| **GSEA-CC** | GO:0000313 | organellar ribosome | 1.27E-07 | 1.09E-07 |
| **GSEA-CC** | GO:0005761 | mitochondrial ribosome | 1.27E-07 | 1.09E-07 |
| **GSEA-CC** | GO:0005740 | mitochondrial envelope | 4.84E-07 | 4.15E-07 |
| **GSEA-CC** | GO:0005747 | mitochondrial respiratory chain complex I | 9.36E-07 | 8.03E-07 |
| **GSEA-CC** | GO:0030964 | NADH dehydrogenase complex | 9.36E-07 | 8.03E-07 |
| **GSEA-CC** | GO:0045271 | respiratory chain complex I | 9.36E-07 | 8.03E-07 |
| **GSEA-CC** | GO:0031966 | mitochondrial membrane | 9.36E-07 | 8.03E-07 |
| **GSEA-CC** | GO:0000315 | organellar large ribosomal subunit | 1.97E-06 | 1.69E-06 |
| **GSEA-CC** | GO:0005762 | mitochondrial large ribosomal subunit | 1.97E-06 | 1.69E-06 |
| **GSEA-CC** | GO:1990204 | oxidoreductase complex | 8.55E-06 | 7.34E-06 |
| **GSEA-CC** | GO:0070069 | cytochrome complex | 1.29E-05 | 1.1E-05 |
| **GSEA-CC** | GO:0022626 | cytosolic ribosome | 2.16E-05 | 1.85E-05 |
| **GSEA-CC** | GO:0015935 | small ribosomal subunit | 4.6E-05 | 3.95E-05 |
| **GSEA-CC** | GO:0005753 | mitochondrial proton-transporting ATP synthase complex | 4.6E-05 | 3.95E-05 |

| | | | | |
|---|---|---|---|---|
| **GSEA-CC** | GO:0045259 | proton-transporting ATP synthase complex | 4.6E-05 | 3.95E-05 |
| **GSEA-CC** | GO:0045277 | respiratory chain complex IV | 5.82E-05 | 5E-05 |
| **GSEA-CC** | GO:0005759 | mitochondrial matrix | 6.61E-05 | 5.67E-05 |
| **GSEA-CC** | GO:0022625 | cytosolic large ribosomal subunit | 0.000168 | 0.000144 |
| **GSEA-CC** | GO:0030684 | preribosome | 0.001214 | 0.001042 |
| **GSEA-CC** | GO:0005838 | proteasome regulatory particle | 0.001552 | 0.001332 |
| **GSEA-CC** | GO:0005730 | nucleolus | 0.001831 | 0.001572 |
| **GSEA-CC** | GO:0005861 | troponin complex | 0.001837 | 0.001577 |
| **GSEA-CC** | GO:0032040 | small-subunit processome | 0.002209 | 0.001896 |
| **GSEA-CC** | GO:0000502 | proteasome complex | 0.003769 | 0.003235 |
| **GSEA-CC** | GO:0005751 | mitochondrial respiratory chain complex IV | 0.003873 | 0.003324 |
| **GSEA-CC** | GO:0120114 | Sm-like protein family complex | 0.004116 | 0.003533 |
| **GSEA-CC** | GO:1905369 | endopeptidase complex | 0.004585 | 0.003935 |
| **GSEA-CC** | GO:0022624 | proteasome accessory complex | 0.004688 | 0.004023 |
| **GSEA-CC** | GO:0000314 | organellar small ribosomal subunit | 0.005035 | 0.004321 |
| **GSEA-CC** | GO:0005763 | mitochondrial small ribosomal subunit | 0.005035 | 0.004321 |
| **GSEA-CC** | GO:0045263 | proton-transporting ATP synthase complex, coupling factor F(o) | 0.005916 | 0.005078 |
| **GSEA-CC** | GO:0097526 | spliceosomal tri-snRNP complex | 0.006168 | 0.005294 |
| **GSEA-CC** | GO:0005681 | spliceosomal complex | 0.01041 | 0.008935 |
| **GSEA-CC** | GO:0005686 | U2 snRNP | 0.010414 | 0.008938 |
| **GSEA-CC** | GO:0005882 | intermediate filament | 0.011298 | 0.009697 |
| **GSEA-CC** | GO:1905368 | peptidase complex | 0.011298 | 0.009697 |
| **GSEA-CC** | GO:0097525 | spliceosomal snRNP complex | 0.01199 | 0.010291 |
| **GSEA-CC** | GO:0045111 | intermediate filament cytoskeleton | 0.01366 | 0.011724 |
| **GSEA-CC** | GO:0016469 | proton-transporting two-sector ATPase complex | 0.013909 | 0.011938 |
| **GSEA-CC** | GO:0030532 | small nuclear ribonucleoprotein complex | 0.014843 | 0.01274 |
| **GSEA-CC** | GO:0005865 | striated muscle thin filament | 0.018497 | 0.015875 |
| **GSEA-CC** | GO:0036379 | myofilament | 0.018497 | 0.015875 |
| **GSEA-CC** | GO:0046930 | pore complex | 0.018497 | 0.015875 |
| **GSEA-CC** | GO:0030017 | sarcomere | 0.025262 | 0.021681 |
| **GSEA-CC** | GO:0034706 | sodium channel complex | 0.02619 | 0.022478 |

| | | | | |
|---|---|---|---|---|
| **GSEA-CC** | GO:0071011 | precatalytic spliceosome | 0.027558 | 0.023652 |
| **GSEA-CC** | GO:0034719 | SMN-Sm protein complex | 0.030175 | 0.025898 |
| **GSEA-CC** | GO:0005839 | proteasome core complex | 0.047326 | 0.040618 |
| **GSEA-MF** | GO:0003735 | structural constituent of ribosome | 4.19E-08 | 3.7E-08 |
| **GSEA-MF** | GO:0005198 | structural molecule activity | 4.19E-08 | 3.7E-08 |
| **GSEA-MF** | GO:0019843 | rRNA binding | 0.003283 | 0.002899 |
| **GSEA-MF** | GO:0030695 | GTPase regulator activity | 0.003283 | 0.002899 |
| **GSEA-MF** | GO:0060589 | nucleoside-triphosphatase regulator activity | 0.003283 | 0.002899 |
| **GSEA-MF** | GO:0008528 | G protein-coupled peptide receptor activity | 0.00338 | 0.002985 |
| **GSEA-MF** | GO:0001653 | peptide receptor activity | 0.005313 | 0.004692 |
| **GSEA-MF** | GO:0016757 | glycosyltransferase activity | 0.016809 | 0.014843 |
| **GSEA-MF** | GO:0005096 | GTPase activator activity | 0.029185 | 0.025772 |
| **GSEA-MF** | GO:0005319 | lipid transporter activity | 0.033882 | 0.029919 |
| **GSEA-MF** | GO:0031492 | nucleosomal DNA binding | 0.034858 | 0.030781 |
| **GSEA-MF** | GO:0005344 | oxygen carrier activity | 0.034858 | 0.030781 |
| **GSEA-MF** | GO:0019825 | oxygen binding | 0.034858 | 0.030781 |
| **GSEA-MF** | GO:0001664 | G protein-coupled receptor binding | 0.034858 | 0.030781 |
| **GSEA-MF** | GO:1901702 | salt transmembrane transporter activity | 0.034858 | 0.030781 |
| **GSEA-MF** | GO:0004497 | monooxygenase activity | 0.038665 | 0.034143 |
| **GSEA-MF** | GO:0016814 | hydrolase activity, acting on carbon-nitrogen (but not peptide) bonds, in cyclic amidines | 0.044492 | 0.039289 |
| **KEGG** | dre00670 | One carbon pool by folate | 0.048291 | 0.038125 |
| **KEGG** | dre03320 | PPAR signalling pathway | 0.049428 | 0.010406 |
| **KEGG** | dre04137 | Mitophagy - animal | 0.049428 | 0.010406 |
| **KEGG** | dre04120 | Ubiquitin mediated proteolysis | 0.049428 | 0.010406 |

**Supplementary Table 7 | Functional enrichment analysis for 5 dpf (MZ*rlig1* vs. WT).** Listed are significantly (padj < 0.05) enriched terms from Gene Ontology (GO) subdivided into biological process (BP), cellular component (CC), and molecular function (MF), gene set enrichment analysis (GSEA) terms, and KEGG pathways.

| Term | ID | Description | padj | q-value |
|---|---|---|---|---|
| **GO-BP** | GO:0051923 | sulfation | 0.033752983 | 0.032168562 |
| **GO-MF** | GO:0020037 | heme binding | 0.039179466 | 0.031807204 |
| **GO-MF** | GO:0046906 | tetrapyrrole binding | 0.039179466 | 0.031807204 |
| **GO-MF** | GO:0031386 | protein tag activity | 0.039179466 | 0.031807204 |
| **GO-MF** | GO:0141047 | molecular tag activity | 0.039179466 | 0.031807204 |
| **GO-MF** | GO:0016712 | oxidoreductase activity, acting on paired donors, with incorporation or reduction of molecular oxygen, reduced flavin or flavoprotein as one donor, and incorporation of one atom of oxygen | 0.039179466 | 0.031807204 |
| **GO-MF** | GO:0031492 | nucleosomal DNA binding | 0.039179466 | 0.031807204 |
| **GO-MF** | GO:0140104 | molecular carrier activity | 0.039179466 | 0.031807204 |
| **GO-MF** | GO:0005344 | oxygen carrier activity | 0.039179466 | 0.031807204 |
| **GO-MF** | GO:0019825 | oxygen binding | 0.039179466 | 0.031807204 |
| **GO-MF** | GO:0044389 | ubiquitin-like protein ligase binding | 0.041240526 | 0.033480441 |
| **GO-MF** | GO:0008146 | sulfotransferase activity | 0.041240526 | 0.033480441 |
| **GO-MF** | GO:0004252 | serine-type endopeptidase activity | 0.041240526 | 0.033480441 |
| **GO-MF** | GO:0061134 | peptidase regulator activity | 0.041993288 | 0.034091559 |
| **GO-MF** | GO:0004175 | endopeptidase activity | 0.043056208 | 0.034954473 |
| **GO-MF** | GO:0008236 | serine-type peptidase activity | 0.043056208 | 0.034954473 |
| **GO-MF** | GO:0017171 | serine hydrolase activity | 0.043056208 | 0.034954473 |
| **GO-MF** | GO:0016782 | transferase activity, transferring sulphur-containing groups | 0.047114915 | 0.038249466 |
| **GSEA-BP** | GO:0051923 | sulfation | 0.015979534 | 0.015716262 |
| **GSEA-BP** | GO:0019953 | sexual reproduction | 0.015979534 | 0.015716262 |
| **GSEA-BP** | GO:0043087 | regulation of GTPase activity | 0.029506084 | 0.029019955 |
| **GSEA-BP** | GO:0071345 | cellular response to cytokine stimulus | 0.036305288 | 0.035707139 |
| **GSEA-BP** | GO:0034612 | response to tumor necrosis factor | 0.037220976 | 0.03660774 |
| **GSEA-BP** | GO:0071356 | cellular response to tumor necrosis factor | 0.037220976 | 0.03660774 |
| **GSEA-BP** | GO:0070374 | positive regulation of ERK1 and ERK2 cascade | 0.037220976 | 0.03660774 |

| | | | | |
|---|---|---|---|---|
| **GSEA-BP** | GO:0070372 | regulation of ERK1 and ERK2 cascade | 0.037220976 | 0.03660774 |
| **GSEA-BP** | GO:0070371 | ERK1 and ERK2 cascade | 0.037220976 | 0.03660774 |
| **GSEA-BP** | GO:0022412 | cellular process involved in reproduction in multicellular organism | 0.037220976 | 0.03660774 |
| **GSEA-BP** | GO:0019221 | cytokine-mediated signalling pathway | 0.037220976 | 0.03660774 |
| **GSEA-BP** | GO:0043547 | positive regulation of GTPase activity | 0.037220976 | 0.03660774 |
| **GSEA-BP** | GO:0034097 | response to cytokine | 0.037220976 | 0.03660774 |
| **GSEA-BP** | GO:0022414 | reproductive process | 0.037220976 | 0.03660774 |
| **GSEA-CC** | GO:0022626 | cytosolic ribosome | 1.21217E-05 | 1.1308E-05 |
| **GSEA-CC** | GO:0005840 | ribosome | 1.21217E-05 | 1.1308E-05 |
| **GSEA-CC** | GO:0044391 | ribosomal subunit | 1.95625E-05 | 1.82493E-05 |
| **GSEA-CC** | GO:0015934 | large ribosomal subunit | 0.001375529 | 0.001283191 |
| **GSEA-CC** | GO:0022625 | cytosolic large ribosomal subunit | 0.003985049 | 0.003717537 |
| **GSEA-CC** | GO:0031105 | septin complex | 0.013681818 | 0.012763373 |
| **GSEA-CC** | GO:0032156 | septin cytoskeleton | 0.013681818 | 0.012763373 |
| **GSEA-CC** | GO:0098803 | respiratory chain complex | 0.019783104 | 0.018455086 |
| **GSEA-CC** | GO:0005940 | septin ring | 0.020194003 | 0.018838402 |
| **GSEA-CC** | GO:0070469 | respirasome | 0.020194003 | 0.018838402 |
| **GSEA-CC** | GO:0034703 | cation channel complex | 0.020194003 | 0.018838402 |
| **GSEA-CC** | GO:0005746 | mitochondrial respirasome | 0.023819857 | 0.022220856 |
| **GSEA-CC** | GO:0022627 | cytosolic small ribosomal subunit | 0.024034369 | 0.022420968 |
| **GSEA-CC** | GO:0098798 | mitochondrial protein-containing complex | 0.034618265 | 0.032294379 |
| **GSEA-CC** | GO:0032153 | cell division site | 0.041725994 | 0.038924974 |
| **GSEA-MF** | GO:0003735 | structural constituent of ribosome | 1.85213E-05 | 1.74748E-05 |
| **GSEA-MF** | GO:0030695 | GTPase regulator activity | 0.005059146 | 0.004773283 |
| **GSEA-MF** | GO:0060589 | nucleoside-triphosphatase regulator activity | 0.005059146 | 0.004773283 |
| **GSEA-MF** | GO:0046873 | metal ion transmembrane transporter activity | 0.008419661 | 0.007943915 |
| **GSEA-MF** | GO:0005096 | GTPase activator activity | 0.014472878 | 0.013655101 |
| **GSEA-MF** | GO:0004252 | serine-type endopeptidase activity | 0.034507689 | 0.032557862 |
| **GSEA-MF** | GO:0005104 | fibroblast growth factor receptor binding | 0.04840491 | 0.045669832 |
| **GSEA-MF** | GO:0008236 | serine-type peptidase activity | 0.04840491 | 0.045669832 |
| **GSEA-MF** | GO:0005216 | monoatomic ion channel activity | 0.04840491 | 0.045669832 |

| | | | | |
|---|---|---|---|---|
| **GSEA-MF** | GO:0004674 | protein serine/threonine kinase activity | 0.04840491 | 0.045669832 |
| **KEGG** | dre00480 | Glutathione metabolism | 0.015806418 | 0.012940927 |
| **KEGG** | dre00983 | Drug metabolism - other enzymes | 0.026898209 | 0.022021926 |
| **KEGG** | dre00591 | Linoleic acid metabolism | 0.000723634 | 0.000592449 |
| **KEGG** | dre00590 | Arachidonic acid metabolism | 0.003599666 | 0.002947095 |

**Supplementary Tables provided as Excel files**

**Supplementary Table 8 | Differentially expressed genes (DEGs) identified in the MZ*rlig1* vs. WT comparison, with samples solely grouped by genotype, independent of behavioural data.** Only DEGs with an absolute $\log_2$ fold change ($|\log_2 FC|$) $\geq 1.5$ and an adjusted p-value (padj) $\leq 0.05$ are included. The table contains the following columns: Gene ID, $\log_2 FC$ ($\log_2$ fold change), p-value (raw p-value), padj (Benjamini–Hochberg adjusted p-value), and Gene name (gene symbol).

**Supplementary Table 9 | Differentially expressed genes (DEGs) identified in 4-cell stage embryos (MZ*rlig1* vs. WT).** The table contains the following columns: Gene ID, $\log_2 FC$ ($\log_2$ fold change), p-value (raw p-value), padj (Benjamini–Hochberg adjusted p-value), and Gene name (gene symbol).

**Supplementary Table 10 | Differentially expressed genes (DEGs) identified in sphere stage embryos (MZ*rlig1* vs. WT).** The table contains the following columns: Gene ID, $\log_2 FC$ ($\log_2$ fold change), p-value (raw p-value), padj (Benjamini–Hochberg adjusted p-value), and Gene name (gene symbol).

**Supplementary Table 11 | Differentially expressed genes (DEGs) identified in shield stage embryos (MZ*rlig1* vs. WT).** The table contains the following columns: Gene ID, $\log_2 FC$ ($\log_2$ fold change), p-value (raw p-value), padj (Benjamini–Hochberg adjusted p-value), and Gene name (gene symbol).

**Supplementary Table 12 | Differentially expressed genes (DEGs) identified in bud stage embryos (MZ*rlig1* vs. WT).** The table contains the following columns: Gene ID, $\log_2 FC$ ($\log_2$ fold change), p-value (raw p-value), padj (Benjamini–Hochberg adjusted p-value), and Gene name (gene symbol).

**Supplementary Table 13 | Differentially expressed genes (DEGs) identified in 1 dpf embryos (MZ*rlig1* vs. WT).** The table contains the following columns: Gene ID, $\log_2 FC$ ($\log_2$ fold change), p-value (raw p-value), padj (Benjamini–Hochberg adjusted p-value), and Gene name (gene symbol).

**Supplementary Table 14 | Differentially expressed genes (DEGs) identified in 5 dpf embryos (MZ*rlig1* vs. WT).** The table contains the following columns: Gene ID, $\log_2 FC$ ($\log_2$ fold change), p-value (raw p-value), padj (Benjamini–Hochberg adjusted p-value), and Gene name (gene symbol).

### Supplementary Movie Legends

**Supplementary Movie 1 | Experimental setup for behavioural assays**. Freely swimming zebrafish larvae are positioned in custom-designed circular acrylic dishes (12 cm). Visual grating stimuli are projected onto the dish from below, and a camera tracks the larvae from above. The optomotor response of the larvae is tested, with live tracking performed in real time using custom-written behavioural tracking software.

**Supplementary Movie 2 | Reconstructed trajectories of representative WT and MZ*rlig1* larvae during a closed-loop optomotor response assay.** The upper dish shows the reconstructed trajectory of a representative WT larva (blue), the lower dish the reconstructed trajectory of a representative MZ*rlig1* larva (magenta). The stimulus sequence consists of an initial period without motion (grey area), followed by a leftward motion stimulus, and ending with a second period without motion (grey area). Each dish has a diameter of 12 cm. The movie is shown in real time.

**Supplementary Movie 3 | Neuronal activity imaging in representative WT and MZ*rlig1* larvae expressing GCaMP8.** Imaging was performed in a single plane of the pretectum (upper row) and the anterior hindbrain (lower row). The WT larva is shown on the left and the MZ*rlig1* larva on the right. The displayed activity traces represent averaged responses to leftward and rightward motion stimuli; during the experiment, 0% coherence trials were presented in between. Each trial lasted 50 s, and the sped-up movies show two stimulus periods compressed to 10 s (10× speed). The stimulus icon (red) indicates the timing of stimulus onset and offset and is shown at real speed. Scale bar: 20 µm.
